# Improved biodiversity detection using a large-volume environmental DNA sampler with in situ filtration and implications for marine eDNA sampling strategies

**DOI:** 10.1101/2022.01.12.475892

**Authors:** Annette F. Govindarajan, Luke McCartin, Allan Adams, Elizabeth Allan, Abhimanyu Belani, Rene Francolini, Justin Fujii, Daniel Gomez-Ibañez, Amy Kukulya, Fredrick Marin, Kaitlyn Tradd, Dana R. Yoerger, Jill M. McDermott, Santiago Herrera

## Abstract

Metabarcoding analysis of environmental DNA samples is a promising new tool for marine biodiversity and conservation. Typically, seawater samples are obtained using Niskin bottles and filtered to collect eDNA. However, standard sample volumes are small relative to the scale of the environment, conventional collection strategies are limited, and the filtration process is time consuming. To overcome these limitations, we developed a new large – volume eDNA sampler with in situ filtration, capable of taking up to 12 samples per deployment. We conducted three deployments of our sampler on the robotic vehicle *Mesobot* in the Flower Garden Banks National Marine Sanctuary in the northwestern Gulf of Mexico and collected samples from 20 to 400 m depth. We compared the large volume (∼40 – 60 liters) samples collected by *Mesobot* with small volume (∼2 liters) samples collected using the conventional CTD – mounted Niskin bottle approach. We sequenced the V9 region of 18S rRNA, which detects a broad range of invertebrate taxa, and found that while both methods detected biodiversity changes associated with depth, our large volume samples detected approximately 66% more taxa than the CTD small volume samples. We found that the fraction of the eDNA signal originating from metazoans relative to the total eDNA signal decreased with sampling depth, indicating that larger volume samples may be especially important for detecting metazoans in mesopelagic and deep ocean environments. We also noted substantial variability in biological replicates from both the large volume *Mesobot* and small volume CTD sample sets. Both of the sample sets also identified taxa that the other did not – although the number of unique taxa associated with the *Mesobot* samples was almost four times larger than those from the CTD samples. Large volume eDNA sampling with in situ filtration, particularly when coupled with robotic platforms, has great potential for marine biodiversity surveys, and we discuss practical methodological and sampling considerations for future applications.

## Introduction

Marine ecosystems are facing a host of anthropogenic threats including global warming, ocean acidification, pollution, overfishing, and invasive species. It is critical to assess the impact of these threats on biodiversity (Brito-Morales et al., 2020; Sala et al., 2021; St John et al., 2016; Worm and Lotze, 2021). Metabarcoding analysis of environmental DNA (eDNA) is an important new tool that can efficiently and effectively help to fill this need (Gallego et al., 2020; Gilbey et al., 2021). DNA sequencing of the trace genetic remains of animals found in bulk environmental samples provides detailed information on the taxonomic makeup of marine communities, and leads to important insights on the diversity, distribution, and ecology of community inhabitants (e.g., Sawaya et al., 2018; Jeunen et al., 2019; Closek et al., 2019; Djurhuus et al., 2020; West et al., 2021; Visser et al., 2021). eDNA analyses are being increasingly applied to mid- and deep-water ocean ecosystems (Canals et al., 2021; Easson et al., 2020; Govindarajan et al., 2021; Laroche et al., 2020; Merten et al., 2021), and advances in robotics and sampling technology could improve sampling strategies to these otherwise difficult to reach regions.

### 1.1 Conventional eDNA sampling approaches

For eDNA analyses in mid and deep-water oceanic environments, seawater is conventionally collected using Niskin bottles, which are triggered to collect water samples at a particular water depth and location. Most commonly, the Niskin bottles are mounted on a conductivity temperature depth (CTD) rosette. A vertical profile of samples can be obtained with the CTD rosette at each location across a range of depths (Andruszkiewicz et al., 2017; Easson et al., 2020; Laroche et al., 2020; Govindarajan et al., 2021). Niskin bottles can also be mounted on other platforms, including remotely operated vehicles (ROVs) (Everett and Park, 2018). Upon recovery, the water samples are immediately filtered, and the filters are preserved for subsequent processing back in the laboratory. Niskin bottle sampling, however, has many limitations. The number, size, and deployment mode (e.g., on a CTD rosette) of the bottles is fixed, which confines experimental design. Sample volumes used for eDNA filtration typically range between 1 to 5 liters and are limited by bottle size, competing scientific needs for sample water, and filtration capabilities (e.g., how quickly and how many samples can be filtered). Relative to the vastness of midwater habitats, these eDNA sampling volumes are minute (Govindarajan et al., 2021; Merten et al., 2021); and may be insufficient for obtaining representative eDNA snapshots, given that eDNA distributions appear to be patchy (Andruszkiewicz et al., 2017). However, the issue of optimizing sample volume is relatively poorly understood relative to other eDNA sampling and processing parameters, such as filter type and DNA extraction protocol (Dickie et al., 2018). Additional considerations for conventional eDNA sampling are the need to use a clean work area and sterile procedures during filtration to reduce the possibility of contamination during processing (Ruppert et al., 2019). Furthermore, the handling time involved for processing water samples collected with Niskin bottles can potentially take several hours, during which time the eDNA samples may experience relatively warm temperatures and eDNA in the samples may potentially decay (Goldberg et al., 2016)

### 1.2 New sampling approaches

Integration of water collection with mobile platforms such as autonomous vehicles, combined with in situ filtration, allows for more efficient water sampling and a greater variety of experimental design possibilities than is achievable with Niskin bottle sampling. For example, Yamahara et al. (2019) coupled the Environmental Sample Processor (ESP) with a long-range autonomous underwater vehicle (LRAUV). Using the LRAUV-ESP, they collected 15 ∼ 0.6 – 1 liter water samples for eDNA analysis over the course of two deployments, although their sampler has the potential to collect up to 60 samples per deployment (Yamahara et al., 2019).

However, the ESP sampler requires approximately one hour to filter one liter of water, and so it may be best suited for applications that require small sample volumes. Autonomous approaches with in situ filtration have also been explored for zooplankton sampling (Govindarajan et al., 2015). In this study, the Suspended Particulate Rosette sampler, originally designed for biogeochemical sampling, was fitted with mesh appropriate for invertebrate larval collection and integrated into a REMUS 600 AUV. “SUPR-REMUS” successfully collected barnacle larvae for DNA barcoding from a coastal embayment with complex bathymetry. For deep-sea environments where target species are relatively dilute, Billings et al. (2017) developed a very large volume plankton sampler for the AUV *Sentry*.

For midwater and deep sea eDNA collection, an approach similar to that described above could be taken, using relevant filter types and seawater sample volumes. Recently, a new autonomous vehicle, *Mesobot,* was designed for studying the ocean’s midwater environments (Yoerger et al., 2021). *Mesobot* can operate fully autonomously or with a fiber optic tether and can survey and unobtrusively track slow-moving midwater animals, as well as collect image and sensor data such as conductivity, temperature, depth, dissolved oxygen, fluorometry and optical backscatter.

*Mesobot* includes a number of features to minimize avoidance and attraction while operating, including white and red LED lighting and slow-turning, large diameter thrusters that reduce hydrodynamic disturbances (Yoerger et al., 2021). *Mesobot* also has payload space to accommodate additional instrumentation, such as an eDNA sampler. The combination of *Mesobot*’s ability to track animals while obtaining imagery and sensor data make it a promising and insightful platform for water column eDNA sampling.

### 1.3 Goals

Our goals were to develop and present a new large-volume autonomous eDNA sampler with in situ filtration mounted on the midwater robot *Mesobot* and assess its utility for conducting midwater eDNA surveys relative to conventional CTD-mounted Niskin bottle sampling. Our study region was the Northwest Gulf of Mexico, and included two sites: Bright Bank in the Flower Garden Banks National Marine Sanctuary, and a deeper water location on the slope of the shelf south of Bright Bank. We sampled at depths ranging from 20 m to 400 m with both methods for their direct comparison. We tested the hypothesis that, because of the larger sample volumes, our eDNA sampler on *Mesobot* (“*Mesobot”* samples) would capture greater animal taxonomic diversity than the CTD – mounted Niskin bottle sampling (“CTD” samples) due to the detection of rare or patchily distributed taxa that were not captured in the small-volume CTD samples. We predicted that taxa identified from the CTD samples would be a subset of those detected in the *Mesobot* samples. As we expected that the most abundant taxa would be present in both sample sets, we also hypothesized that despite the differences in taxon detection, that overall patterns of community structure identified by the two approaches would be similar. To test these hypotheses, we sequenced the V9 barcode region of 18S rRNA to analyze the metazoan eDNA community and compared biodiversity metrics from both sample types. We also described the utility of our eDNA sampler for marine midwater biodiversity surveys, focusing on the topics of sampling volume and practical methodological issues.

## 2 Material and Methods

### 2.1 Study site

We conducted a cruise on the *R/V Manta* in September of 2019 out of Galveston, Texas, USA. The CTD samples presented here are a subset of a larger regional survey. Our focal site was Bright Bank, located in the Northwest Gulf of Mexico off of the coasts of Louisiana and Texas (Fig. 1). Bright Bank received federal protection in March 2021 as part of the recent expansion of the Flower Garden Banks National Marine Sanctuary (FGBNMS). Bright Bank is a shelf-edge carbonate bank that hosts a diverse mesophotic reef ecosystem spanning 117 to 34 m depth (https://flowergarden.noaa.gov/) and is an important habitat for commercially-important and threatened fish species (Dennis and Bright, 1988; Sammarco et al., 2016). We sampled eDNA using both the *Mesobot* sampler and CTD casts at two sites: 1) “Bright Bank” site, located within 3 nautical miles of the center of the bank; and 2) “Slope” site located in offshore water at the slope of the continental shelf, approximately 21 nautical miles south of the bank and with a water depth of approximately 500 m. No permits were required for our work.

**Fig. 1.**
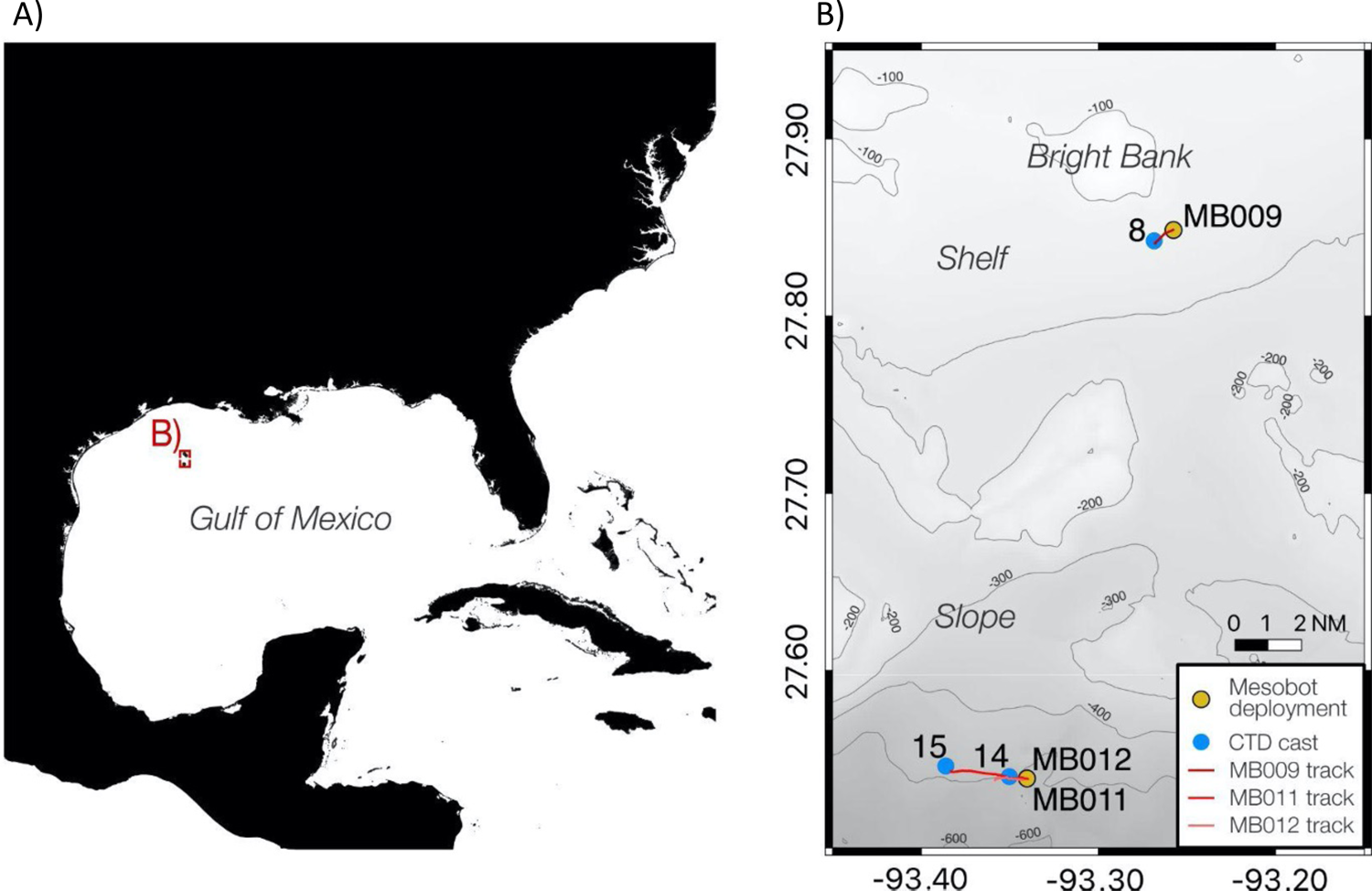
Map of study area. A) location in the Gulf of Mexico; B) close up of study area including Bright Bank and the deeper site. Blue dots indicate CTD locations and yellow dots indicate *Mesobot* deployment locations (MB009, MBOl 1, and MB012). Red lines indicate the *Mesobot* tracks.

### 2.2 Large-volume eDNA sampler with in situ filtration

We developed an adjustable volume eDNA sampler capable of filtering large seawater volumes (10s to 100s of liters) that can be mounted on autonomous platforms such as the hybrid robotic vehicle *Mesobot* (Fig. 2; Fig 3; Supplementary Fig. 1). The eDNA sampler consists of 12 pumps and 12 filters with one pump per filter. The sampler includes two identical pump arrays, originally designed and built as the core of the Midwater Oil Sampler (MOS), an AUV water sampler for oil spills. Each MOS pump array contains six submersible pumps (Shenzhen Century Zhongke Technology model DC40-1250) and a microprocessor that enables an external computer to command individual pumps and log pump status through an RS232 serial connection. The MOS pump array is potted in polyurethane and pressure tested to 6000 m depth. Water enters each filter-pump pair through a unique intake tube. After passing through the pump, the water exits the assembly through a common discharge tube where a flowmeter (Omega Engineering FPR-301) measures the flow. Flow measurements are processed and communicated to *Mesobot* at a frequency of 10 Hz by a secondary microrprocessor mounted inside *Mesobot’s* main housing. We built two spare pump arrays, so that upon retrieval of *Mesobot*, the used sampler can be quickly exchanged with a clean sampler.

**Fig. 2.**
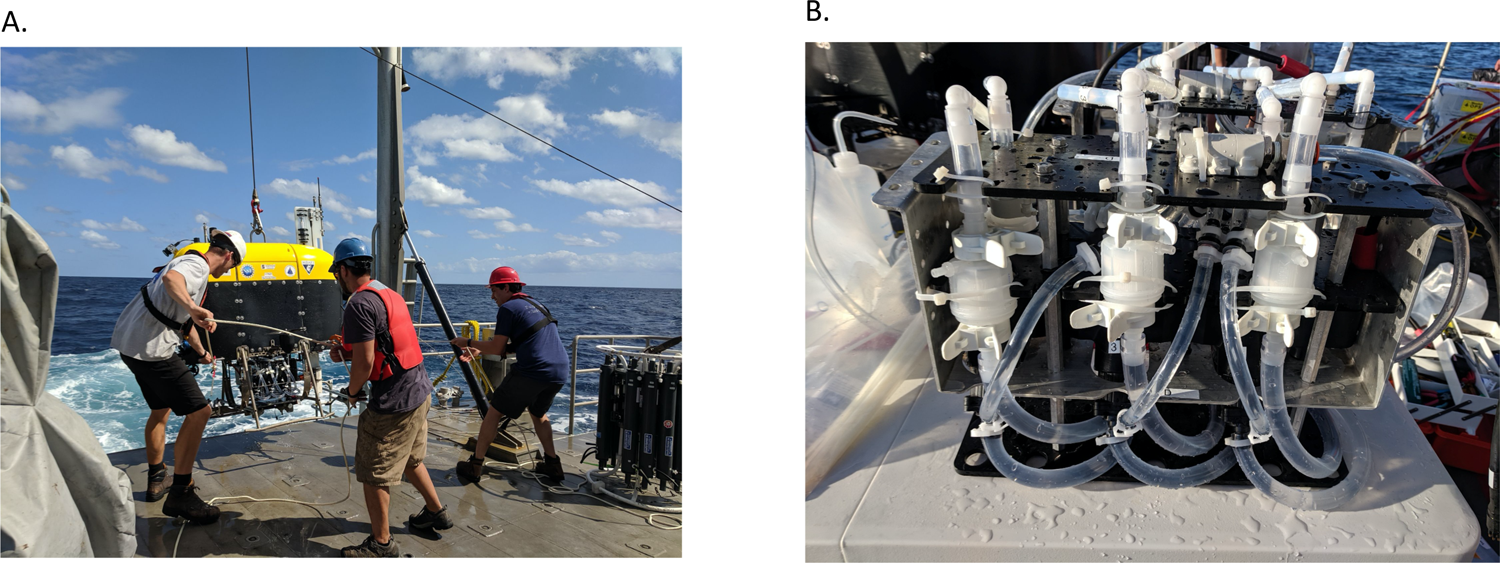
A) *Mesobot* with the eDNA sampler being retrieved after a deployment on the R/V Manta; B) close-up of the eDNA sampler.

**Fig. 3.**
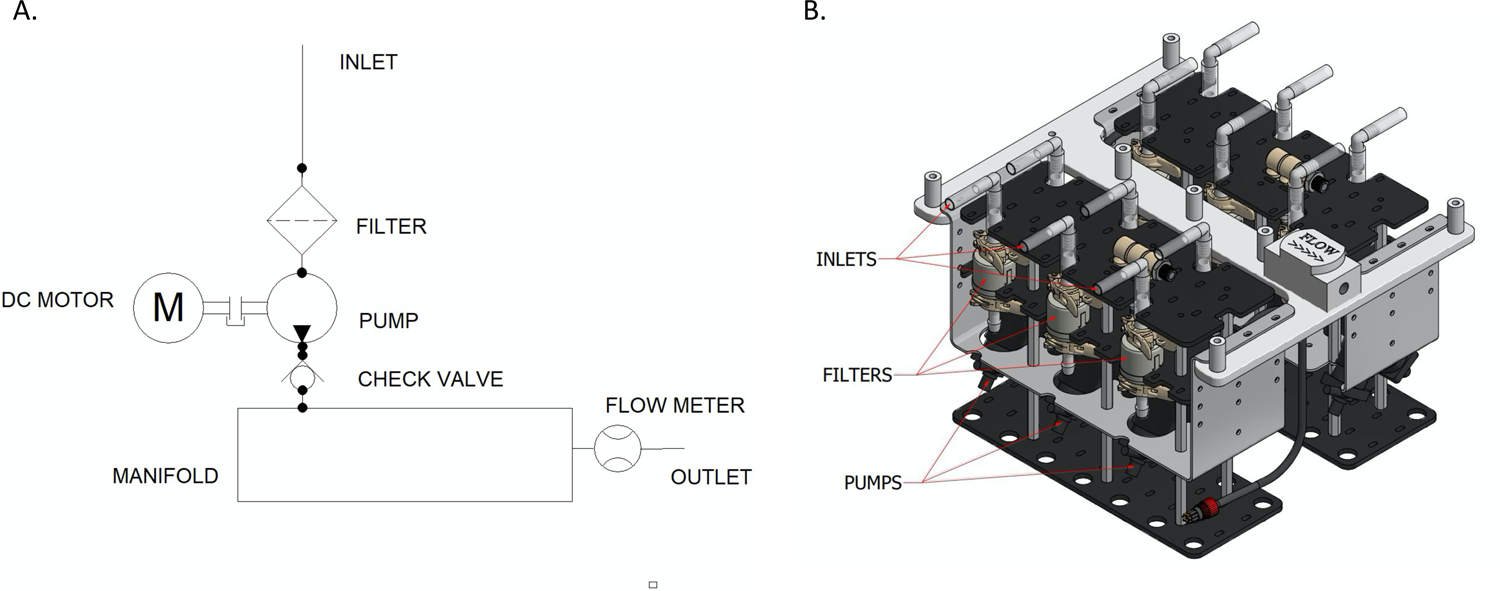
Sampler design. A) Schematic of one pump/filter channel. Each sampler has 6 such channels that flow into a common manifold with an outlet through a single flowmeter. All 6 pumps are controlled by a single microcontroller; B) CAD drawing of the complete sampler. *Mesobot* carried two such samplers for a total of 12 pump/filter units on each dive.

The pumps are connected by bleach-sterilized plastic tubing to Mini Kleenpak capsule filters (Pall Corporation, Port Washington, New York, USA; cat. # KA02EAVP8G). Each filter is individually encapsulated and consists of an inner 0.2 µm Polyethersulfone (PES) filter and an outer PES pre-filter with a variable pore size, resulting in an effective filtration area of 200 cm^2^ for the entire filter capsule. Check valves prevent backflow from reaching any of the filters. Each pump filters seawater at a rate of approximately 2 L/min. Only one pump per MOS pump array can be run at a time, but both arrays can be run simultaneously allowing for duplicate samples to be taken at each of six sampling events.

The eDNA sampler was mounted on the underside of *Mesobot* (Fig. 2). The timing and duration of sampling events were controlled by the main control computer inside the main housing of the *Mesobot* and communicated to the sampler via the secondary microprocessor. To ensure that samples were taken at the proper time, pump commands were interleaved in the mission control program sequence which includes motion commands such as depth changes.

### 2.3 Sampler deployments on Mesobot

Three fully autonomous, untethered *Mesobot* dives were conducted at the Bright Bank (dive MB009) and the Slope (dives MB011 and MB012) sites (Table 1). Prior to each dive, the sampler tubing was cleaned with 10% bleach and rinsed multiple times with ultrapure water. The sampler pumps were then primed by filling the filter capsules with ultrapure water. All filters had been sterilized by autoclaving before the cruise. An additional sealed filter capsule that was filled with ultrapure water was attached to *Mesobot*’s base to serve as a field control. It took approximately an hour and a half of time to complete the pre-dive sampler cleaning and priming steps by one person. At the start of each dive, *Mesobot* was lowered into the water from the vessel’s A-frame and then released. *Mesobot* then executed the programmed sequence of depth changes and sampling operations. During these dives, *Mesobot* used its control system and thrusters to hold depth precisely (+/- 1cm) while drifting with the ambient currents, much like a Lagrangian float. During *Mesobot* deployments, an acoustic ultra-short baseline (LinkQuest TrackLink) tracking system was used to determine the position and depth of the AUV underwater. During each dive, *Mesobot* could drift several kilometers, accordingly we used the tracking information to follow the vehicle as it drifted and to ensure that the vessel was positioned appropriately to recover the vehicle when it returned to the surface at the end of the dive. To help locate the vehicle after it surfaced, the vehicle carried 3 strobe lights, a VHF beacon, and an Iridium/GPS unit that transmitted the vehicle’s surface position through a satellite link. The additional surface recovery aids were important on the last dive, MB012, when the USBL tracking system failed and the vehicle surfaced at night time about a kilometer from the expected position.

**Table 1.** Summary of samples collected, including the *Mesobot*-mounted sampler samples and the CTD-mounted Niskin bottle samples. Additional sampling details for the *Mesobot* samples are in Supplementary Table 1 and details for the CTD samples are in Supplementary Table 2.

| Cast or Dive | Date | Time (UTC) | Site | Station | Latitude | Longitude | Depth range (m) | # samples |
| --- | --- | --- | --- | --- | --- | --- | --- | --- |
| 8 | 9/25/19 | 16:29 | Bright Bank | Bright Bank | 27.84239 | -93.268503 | 100 - 40 | 11 |
| 14 | 9/26/19 | 17:36 | Slope | Slope | 27.54012 | -93.35027 | 100 - 40 | 12 |
| 15 | 9/26/19 | 21:01 | Slope | Slope | 27.54607 | -93.38611 | 400 - 160 | 11 |
| MB009 | 9/25/19 | 15:25 | Bright Bank | Bright Bank | 27.8485 | -93.2576 | 20 - 120 | 12 |
| MB011 | 9/26/19 | 17:11 | Slope | Slope | 27.53905 | -93.34029 | 200 - 400 | 12 |
| MB012 | 9/26/19 | 23:29 | Slope | Slope | 27.53905 | -93.34029 | 40 - 320 | 12 |

For all deployments, twelve samples (consisting of 6 sets of duplicates, which served as biological replicates) were collected along vertical transects. At the Bright Bank site, samples were taken between 120 and 20 m; at the Slope site, samples were taken between 400 and 40 m over the course of two deployments (Table 1). Once *Mesobot* was recovered after each deployment, the filter capsules were removed from the sampler and drained, and the ends were sealed with parafilm. The sealed filter capsules were stored in coolers filled with dry ice within a few minutes of retrieval.

### 2.4 Conventional CTD – mounted Niskin bottle sampling

Seawater samples were collected using a Seabird SBE 19 CTD rosette equipped with twelve 2.5-liter Niskin bottles. Samples were collected in triplicate (i.e., three biological replicates) at four depths in each cast, with the target depths selected to complement the *Mesobot* sampling depths (Table 1). At the Bright Bank site, one CTD cast (“Cast 8”) was conducted and samples were collected between 40 and 100 m depth. At the slope site, two CTD casts were conducted and samples were collected at depths ranging from 40 to 100 m (“Cast 14”) and from 160 to 400 m (“Cast 15”) (Table 1).

Once on board the ship, seawater from each Niskin bottle was either transferred to a sterile Whirl-Pak stand-up sample bag (Nasco Sampling, Madison, WI, USA) and filtered in the wet lab, or directly filtered from the Niskin bottle on deck. The entire volume of seawater from each bottle was filtered through a sterile 0.22 μm PES Sterivex filter (MilliporeSigma, Burlington, MA USA). Sterivex filters have a surface area of 10 cm^2^. Water was filtered using a Masterflex L/S peristaltic pump (Masterflex, Vernon Hills, IL, USA) set to 60 RPM equipped with four Masterflex Easy-load II pump heads using Masterflex L/S 15 high-performance precision tubing. Prior to each cast, the tubing was sterilized by pumping a 10% bleach solution for 5 minutes with the pump set at 60 RPM. The tubing interior was then rinsed thoroughly by pumping ultrapure water for 5 minutes at the same flow rate. Following sample filtration, residual water was pumped out of the Sterivex filters, the filters were placed in sterile Whirl-pak bags, and the bags were placed on dry ice in a cooler for the remainder of the cruise. The volume of filtered water was measured with a graduated cylinder and recorded. The average volume of water filtered per Niskin bottle was 2.22 ± 0.25 SD liters. For each CTD cast, a field control consisting of approximately 2 liters of ultrapure water was also processed in the same manner and using the same equipment as the field samples. The total shipboard processing time for the Niskin bottles was approximately two hours per cast with two people. Upon return to port in Galveston, TX, the CTD and the *Mesobot* samples were shipped on dry ice to Woods Hole, MA. Upon arrival in Woods Hole, the filters were stored in a −80°C freezer until DNA extraction, which took place approximately three months later.

### 2.5 eDNA extraction

For the *Mesobot* samples, Mini Kleenpak capsules were opened using a UV-sterilized 3-inch pipe cutter and the outer and inner PES filters were removed and dissected from the capsules using a sterile scalpel and forceps. Each inner and outer filter was cut into six pieces, which were placed into sterile 5 ml centrifuge tubes, and the DNA was extracted from each of the 12 fractions of the filter using DNEasy Blood & Tissue DNA extraction kits (Qiagen, Germantown, MD, USA), with some modifications to the protocol. 900 ul of Buffer ATL and 100 ul of proteinase K were added to each 5 ml centrifuge tube. The tubes were incubated at 56° for 3 hours and vortexed periodically during the incubation period. Following the incubation, 1000 μL of buffer AL and ethanol were added to each centrifuge tube. The entire volume of the lysate was spun through a single spin column in five steps. Washes were performed according to the manufacturer’s protocol, and DNA extracted from each filter piece was eluted in 80 μL of AE buffer. The inner and outer filters for each 1/6^th^ portion were extracted separately, resulting in a total of 12 extractions per sample. The DNA concentration of each filter piece extraction was measured with a Qubit fluorometer (Life Technologies, Carlsbad, CA, USA) using the 1X High-sensitivity double-stranded DNA assay. Equal volumes of all inner 1/6^th^ fractions were pooled yielding a pooled DNA extract for the inner filter for each sample. Outer 1/6^th^ fractions were pooled in the same manner, resulting in a pooled DNA extract for the outer filter for each sample. These two pooled DNA extracts were processed separately for subsequent PCR, library preparation and sequencing.

For the CTD samples, genomic DNA from the Sterivex filters was extracted using DNEasy Blood & Tissue extraction kits following the manufacturer’s protocol adapted to accommodate the Sterivex filter capsules (Govindarajan et al., 2021). DNA was eluted in 80 μL of molecular-grade water. The DNA concentration of each Sterivex filter extraction was also measured with the Qubit 1X High-sensitivity double-stranded DNA assay.

### 2.6 Library preparation and sequencing

Library preparation and sequencing followed the approach in Govindarajan et al. (2021) with a few modifications. All PCR samples were diluted 1:10 in molecular-grade water to prevent possible inhibition (Andruszkiewicz et al., 2017). Duplicate 2.5 µl aliquots from each sample were amplified in 25 μL reactions with 12.5 μL of KAPA HiFi HotStart ReadyMix (Kapa Biosciences, Wilmington, MA, USA), 0.5 μL of 10 μM forward and reverse primers (final concentrations of 0.200 μM), and 9 μL of molecular-grade water. The primers used were 1380F and 1510R, which amplify the V9 portion of the 18S rRNA gene (Amaral-Zettler et al., 2009) with CS1 and CS2 linkers for subsequent ligation of Fluidigm adaptors. The primer sequences with linkers are: ACACTGACGACATGGTTCTACACCCTGCCHTTTGTACACAC (1380F-w-CS1-F) and TACGGTAGCAGAGACTTGGTCTCCTTCYGCAGGTTCACCTAC (1510R-w-CS2-R). Primers were ordered from Eurofins Genomics (Louisville, KY, USA) at 100 μM concentration in TE buffer and diluted to 10 μM to prepare the PCR reactions. Cycling conditions included an initial denaturation step at 95°C for 3 minutes; 25 cycles of 95°C for 30 seconds, 55°C for 30 seconds, and 72°C for 30 seconds; and a final extension step of 72°C for 5 minutes. PCR products were visualized on a 1% agarose gel in TBE buffer stained with GelRed (Biotium, Fremont, California, USA) to determine the presence of amplicons of the expected size. The duplicate PCRs were pooled and sent to the Genome Research Core at the University of Illinois at Chicago (UIC).

At the UIC Genome Research Core, a second round of PCR amplification was conducted to ligate unique 10-base barcodes to each PCR product. The PCR was conducted using MyTaq HS 2X master mix and the Access Array Barcode Library for Illumina (Fluidigm, South San Francisco, CA, USA). Cycling conditions included an initial denaturation step at 95°C for 5 minutes; 8 cycles of 95°C for 30 seconds, 60°C for 30 seconds, and 72°C for 30 seconds; and a final 7-minute extension at 72°C. The barcoded PCR products were pooled and purified using 1.0X Ampure beads (Beckman Coulter, Indianapolis, IN, USA). This method retains amplicons (with primers, linkers, and adapters) longer than 200 bp.

An initial paired-end, 150-basepair sequencing run on an Illumina MiniSeq platform was conducted to determine the expected number of reads per sample. Equal volumes of each library were pooled, and the pooled libraries with a 15% phiX spike-in were sequenced. The volumes of each sample to be pooled for subsequent sequencing on an Illumina MiSeq were adjusted based on the relative number of reads produced by the initial MiniSeq run. Our goal was to obtain an equal sequencing depth among all field samples. Volumes pooled ranged from 1.0 to 30.0 μL. The vast majority of the negative controls (filtration blanks, extraction blanks, and no-template controls) produced very few reads on the MiniSeq run. One μL of each was pooled to increase the overall sequencing effort of the field samples; however, for the *Mesobot* filtration blanks, the volume was adjusted in the same manner as for the field samples. The volume-adjusted libraries were loaded on to a MiSeq platform and sequenced using v2 chemistry targeting paired-end 250 bp reads. De-multiplexing of reads was performed on the instrument. In addition to our sampler and Niskin bottle samples, additional Niskin bottle samples from the larger Bright Bank survey and their associated controls were also included on the sequencing runs. As these samples and controls were processed along with our focal samples, we included these additional controls in our sequence quality control (described below). In total, three MiSeq runs were conducted with the intent of obtaining a target depth of approximately 100,000 reads per sample.

### 2.7 Contamination controls

Rigorous procedures to prevent and monitor contamination were taken at every step from sample collection through sequencing. During sampling filtration, all surfaces in the wet lab were cleaned with 10% bleach and rinsed multiple times with ultrapure water before every use. Nitrile gloves were worn and changed often. Field controls were taken for every *Mesobot* and CTD sampling event as described above. Back on shore, DNA extractions were conducted at WHOI in the Govindarajan lab and PCR reactions were prepared at Lehigh University in the Herrera lab.

Post-PCR products were handled for gel electrophoresis in a separate laboratory space at Lehigh University. All procedures in the WHOI, Lehigh, and UIC sequencing laboratories included the following measures to ensure sample integrity: 1) Nitrile lab gloves were always worn and changed frequently; 2) Pipettes were UV-sterilized before use and sterile filter tips were used; 3) All lab surfaces were cleaned with 10% bleach and rinsed with Milli Q water before each use; 4) PCR preparations were conducted in a PCR hood with a HEPA filter with positive airflow, and the work space was additionally decontaminated with UV light before each use; 5) Field controls were extracted, amplified and sequenced alongside the field samples; and 6) Six DNA extraction blanks were amplified and sequenced, and two PCR no-template controls (NTC) were included in each plate for the first round of PCR, pooled and sequenced.

None of the negative controls (filtration blanks, extraction blanks and PCR NTCs) produced visible amplicons after the first PCR, and the vast majority produced far fewer sequencing reads than the field samples, as expected (105 ± 137 s.d. vs 33,902 ± 25,543 s.d.). Two of the control sample libraries, a field negative control from a CTD cast not included in the data analysis and a PCR no-template control, produced more reads than expected (12,385 and 5,299, respectively). These and four other samples were re-sequenced to obtain correct data for the misprocessed field control and to validate our initial sequencing results (Appendix 1).

### 2.8 Bioinformatics

Sequencing data was received as demultiplexed fastq.gz files for each sample and was processed using Quantitative Insights Into Microbial Ecology 2 (QIIME2) version 2020.11 (Bolyen et al., 2019), following the general approach described in Govindarajan et al. (2021). Raw data was deposited in Dryad. Sequence quality plots were examined, forward primer sequences at the 5’ end and reverse complements of reverse primers at the 3’ end were trimmed using the Cutadapt QIIME2 plugin (Martin, 2011). Sequences were quality filtered, truncated to 120 base pairs in length, denoised, and merged using DADA2 (Callahan et al., 2016) within the QIIME2 platform. Sequences from each run were processed separately and merged after the DADA2 step.

Singleton and doubleton (summed through the dataset) ASVs were removed from further analysis. These and subsequent merging and filtering steps were accomplished using the QIIME2 feature-table plugin. The resulting amplicon sequence variants (ASVs) were taxonomically classified using a naïve Bayesian classifier (Bokulich et al., 2018) that was trained on the Silva v.132 99% small subunit rRNA database (Quast et al., 2013) for the 18S V9 amplicon region.

For each ASV in the dataset that was present in both the samples and in any of the controls, the maximum number of reads found in any control was subtracted from every sample (0.84% of the sample dataset). An additional 143 reads (0.00086% of the remaining sequences) that were classified as human and insect were removed. The resulting dataset was then filtered to include metazoan sequences only. Sampler inner and outer filters were analyzed both separately and together. Biodiversity was visualized using broad taxonomic categories (Silva levels 6 and 7; generally corresponding to order or family, respectively). The V9 marker is not used for species – level identification and species – level identification was outside the scope of this work. Rarefaction curves were generated in QIIME2 to assess and compare sequencing depths. After randomly sampling the data from each sample to the lowest sequencing depth of any field sample, Bray-Curtis dissimilarities were calculated in QIIME2 and were used to generate non-metric multidimensional scaling (nMDS) plots with sampling depth and sample type (*Mesobot* or CTD) visualized using the package vegan 2.3_5 (Oksanen et al., 2016) in R Version 4.0.4 (R Core Team, 2021). For the *Mesobot* filters, nMDS plots were also generated to compare the diversity collected on inner and outer filters. In this analysis, 4 samples with exceptionally low read counts on the inner filter were excluded, as described in the results section. Functional regressions of sampling depth against each nMDS axis were conducted to assess the significance of observed patterns (Ricker, 1973). Permutational multivariate analysis of variance (PERMANOVA) tests were conducted using the “adonis” function in vegan to assess the effects of sample type, sampling depth, and for *Mesobot* filters, inner and outer filter type. Taxon comparisons between sample categories (e.g., filter type, sampling approach, depth) were performed using an online Venn diagram tool from the University of Ghent (http://bioinformatics.psb.ugent.be/webtools/Venn/).

## 3 Results

### 3.1 Sampler performance, and sample collection summary

The *Mesobot* sampler collected a total of 36 samples on three successful deployments (Table 1; Supplementary Table 1). Duplicate samples at 6 depths were obtained in each deployment, for a total of 12 samples per deployment. In the first deployment (MB009), the sampler pumps ran for 20 minutes at 20 m depth intervals between 120 m and 20 m. In the second deployment (MB011), the sampler took 30-minute samples at 40 m depth intervals between 400 m and 200 m. In the third deployment (MB012), the sampler took one pair of samples filtering for 30 minutes at 320 m, and additional sample pairs filtering for 20 minutes at depths of 160 m, 100 m, 80 m, 60 m, and 40 m. For all deployments, the sampler flow rate was slightly over 2 liters per minute. The flow rate typically declined gradually over the sampling period, consistent with our expectation that material was accumulating on the filters (Fig. 4).

**Fig. 4.**
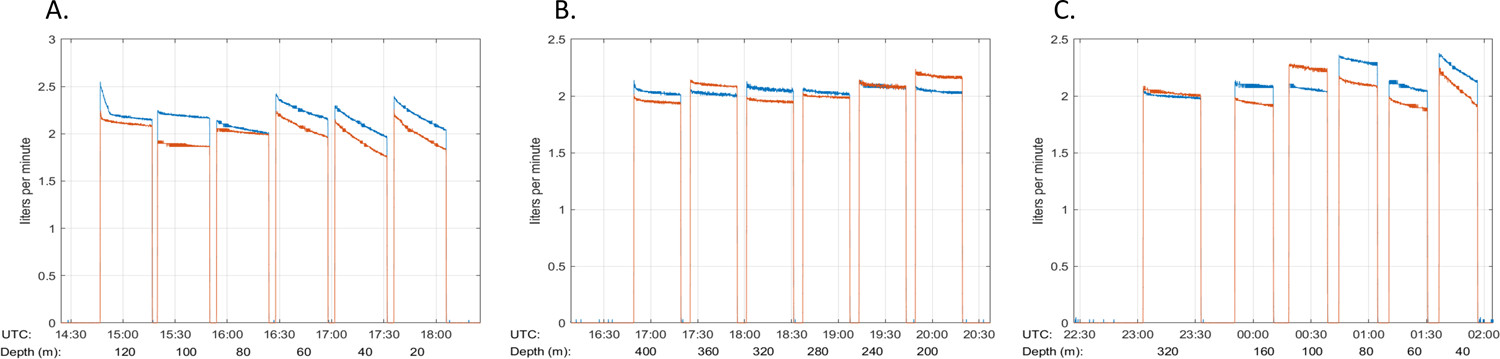
*Mesobot* sampler flow rates over time. The red and blue lines represent the flow rate from duplicate pumps. A) MB009 (Bright Bank site); B) MB011 (Slope site); C) MB012 (Slope site).

### 3.2 CTD data and Niskin bottle sample collection summary

A total of 34 eDNA samples were collected with Niskin bottles over 3 CTD casts (Table 1; Supplementary Table 2). Twelve Niskin bottles were deployed on each CTD cast, but one sample was lost from Cast 8 (100 m) and another from Cast 15 (400 m) due to bottle malfunctions. The CTD profiles from these casts indicated a stratified water column with a thermocline beginning around 40 m at the Bright Bank site and 50 m at the Slope site, with the deep chlorophyll maximum (DCM, corresponding to peak fluorescence) slightly deeper than the thermocline (Supplementary Fig. 2).

### 3.3 Total eDNA yield

As expected given the larger sample volumes, the sampler collected more eDNA than the Niskin bottle sampling. However, the eDNA yield per liter of water filtered was comparable between methods for samples collected at the same depth (Fig. 5). eDNA concentration yields were higher in shallower water (i.e., less than 100 m), with the highest yields (up to ∼8 ng/µl) roughly coinciding with the approximate depth of the DCM (60 m) (Supplementary Fig. 2). eDNA yields were much lower (<1.5 ng/µl) at sampling depths greater than 100 m. For the *Mesobot* samples, the inner filters generally yielded slightly higher (i.e., within a couple ng/µl) DNA concentrations than the outer filters, with greater variation at the Bright Bank site, where one inner filter yielded ∼30 ng/ul more DNA than its corresponding outer filter (Fig. 6). For any given inner or outer filter from a *Mesobot* sample, the DNA concentrations of the extractions stemming from individual filter pieces were relatively similar in most cases, but a few samples (particularly those with the higher overall DNA yields) showed substantial variation (Fig. 6).

**Fig. 5.**
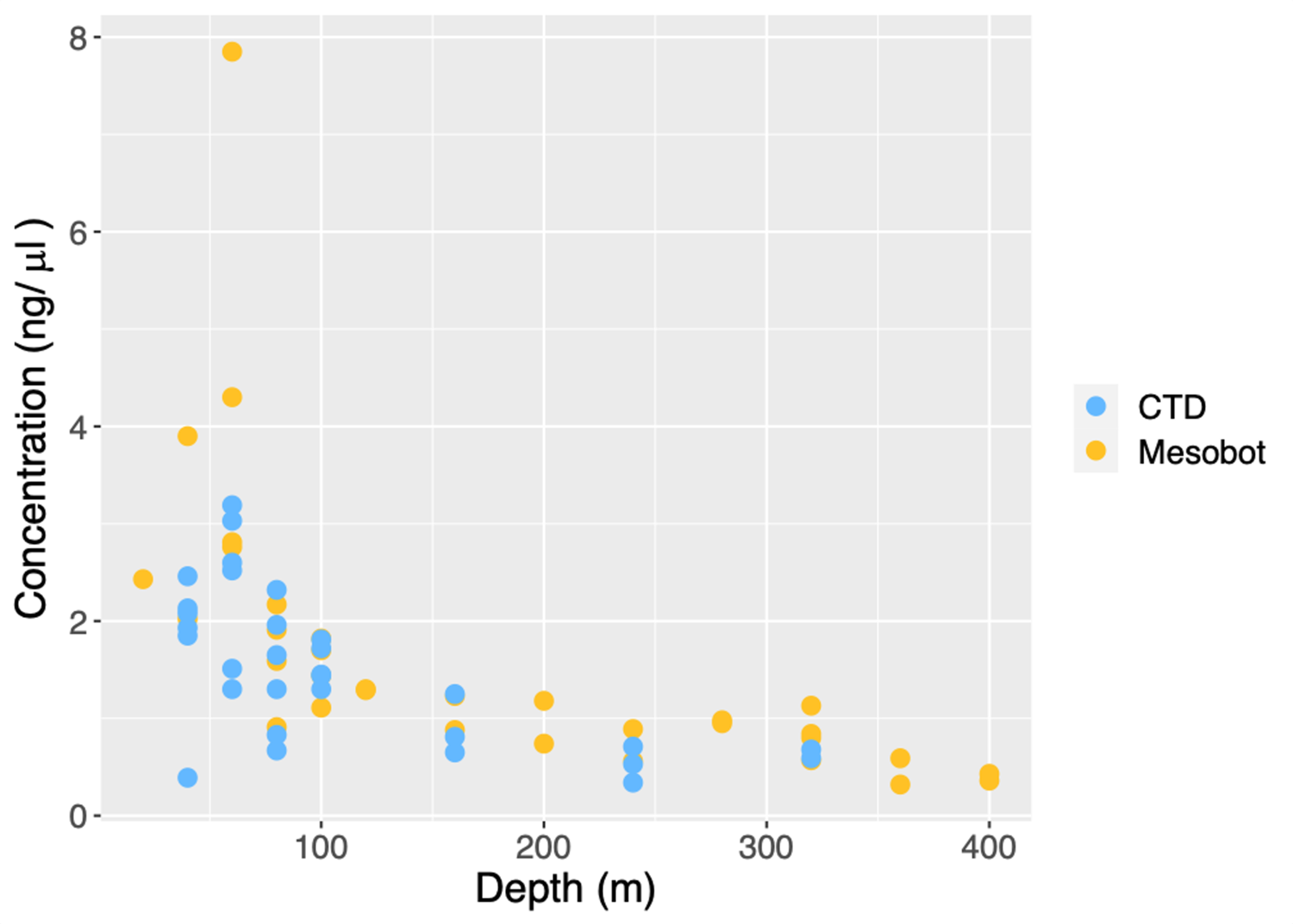
DNA yield (total ng of DNA recovered per liter of water sampled) versus depth for the *Mesobot* and CTD samples. *Mesobot* sample yields are the sum of individually-extracted filter pieces divided by the sample volume. Concentrations from individual inner and outer filter pieces are shown in Fig. 6.

**Fig. 6.**
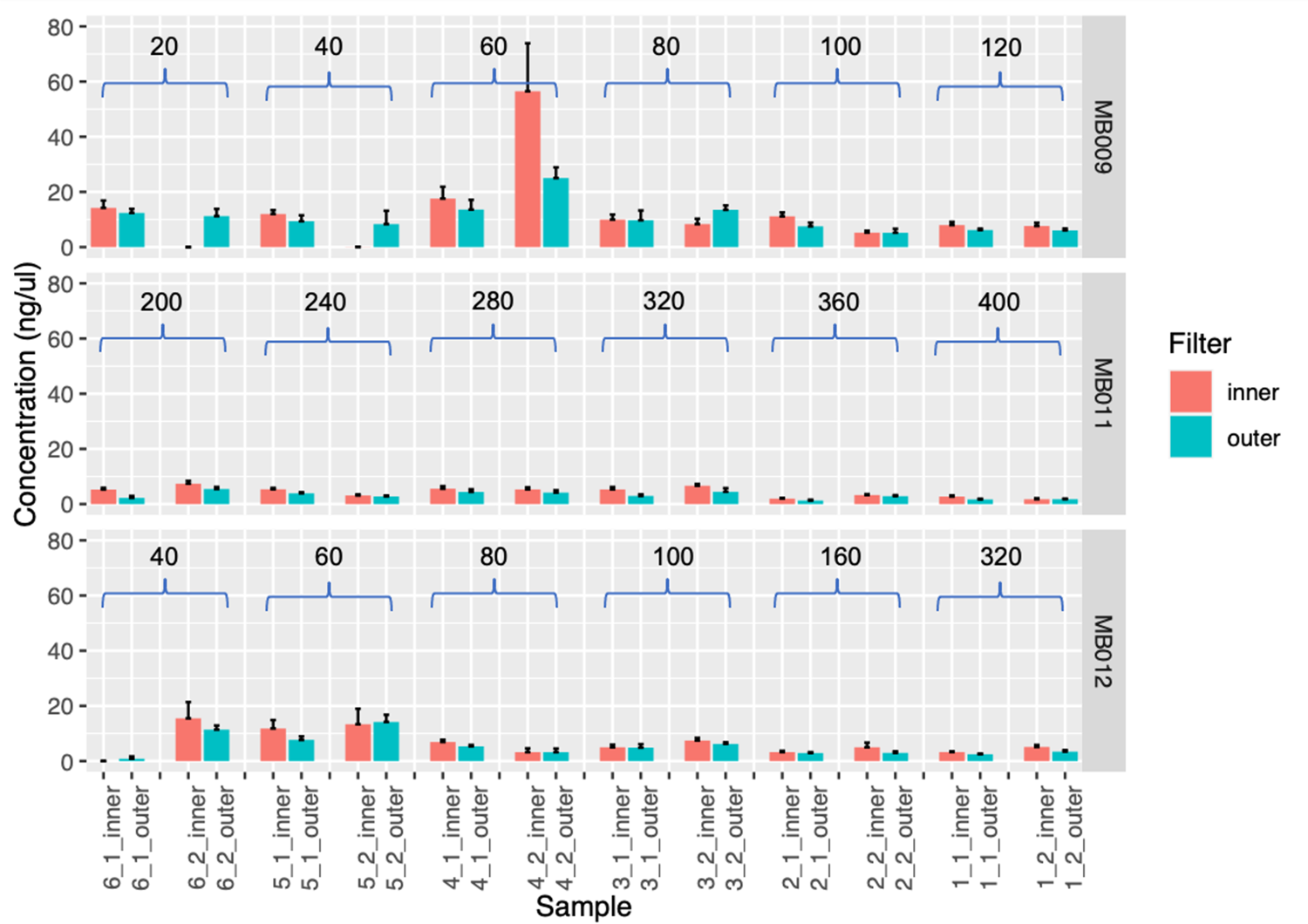
DNA concentrations (mean +/- standard deviation) of inner and outer filter pieces from each *Mesobot* sample. Sampling depth (m) is indicated above bars. MB009 originates from the Bright Bank site and MB011 and MB012 originate from the Slope site.

### 3.3 Metazoan sequence diversity

The number of metazoan reads varied greatly within and between *Mesobot* sampler and CTD datasets, and also between the Mini Kleenpak inner (*Mesobot*-inner, “MBI”) and outer (*Mesobot*-outer; “MBO”) filter dataset (Table 2; Supplementary Table 3). The MBO dataset consisted of 36 samples with 1,096 metazoan ASVs and 2,700,417 metazoan sequences. The mean number of reads per sample ranged from 23,530 to 207,391 with a mean of 75,012. The MBI dataset, with 36 samples, in general had fewer metazoan ASVs (703), total sequences (582,246) and reads per sample (mean = 16,173.5 reads, min = 3 reads; max = 68,149 reads). For a given *Mesobot* sample, the majority of metazoan reads originated from the outer filter, both in terms of the percent of metazoan reads in the dataset (Fig. 7) and in the absolute number of metazoan sequences (Supplementary Table 3). *Mesobot* samples from Bright Bank (MB009) in general had proportionately more metazoan sequences on the outer filter than those from the Slope site (MB011 and MB012) (Fig. 7).

**Fig. 7.**
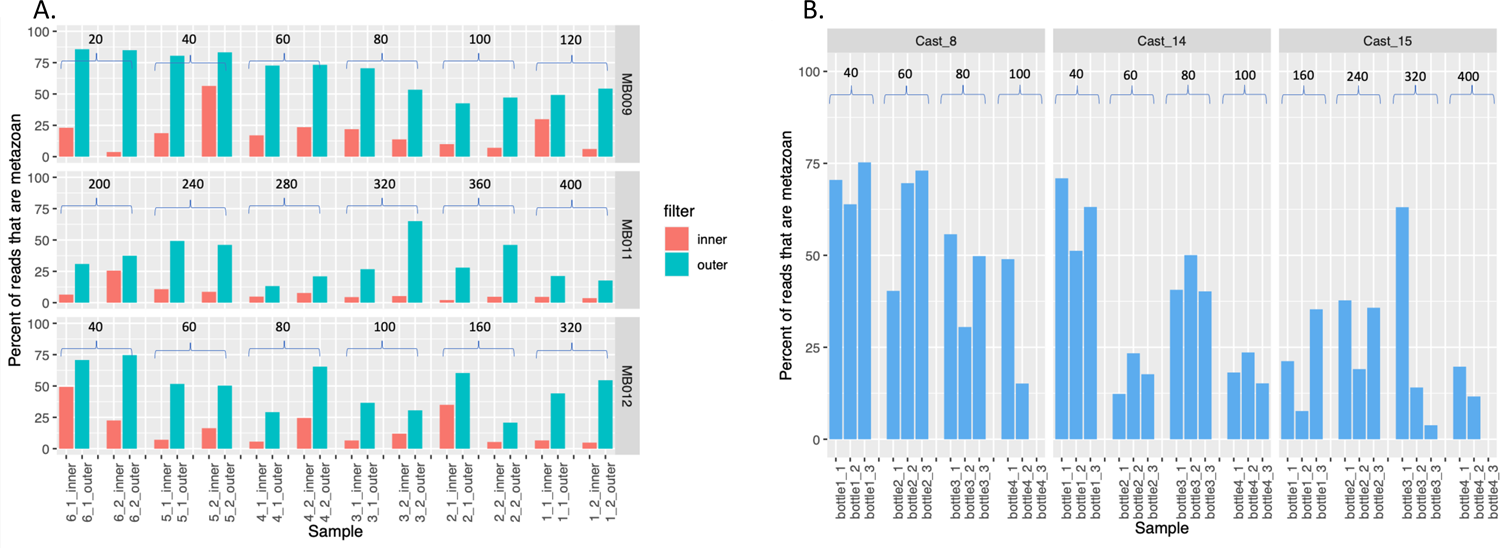
Percent metazoan and non-metazoan reads from the A) inner and outer *Mesobot* sample filters; and B) CTD samples. Sampling depth (m) is indicated above the bars. Note we do not have samples for one of the replicates of Cast 8 - 100 m and for Cast 15 - 400 m, due to bottle mishaps. MB009 and Cast 8 originate from the Bright Bank site and MB011, MB012, Cast 14, and Cast 15 originate from the Slope site.

**Table 2.**
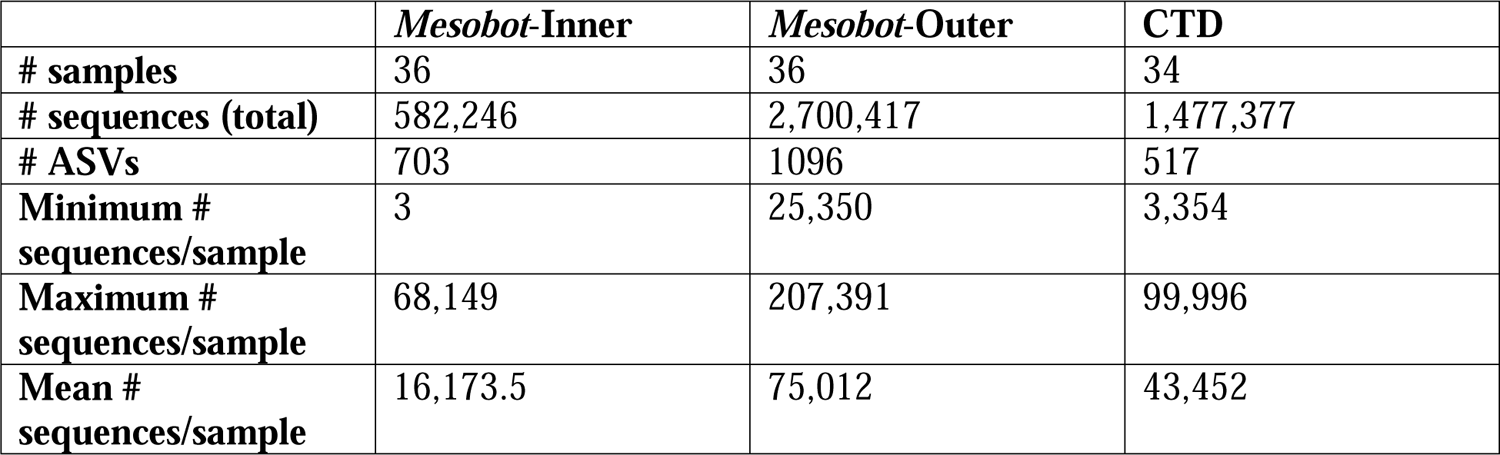
Metazoan sequence summary.

The CTD dataset included 34 samples with 517 metazoan ASVs and 1,477,377 metazoan sequences. The number of metazoan reads per sample ranged from 3,354 to 99,996, with a mean of 43,453, and in most samples, represented less than half of the total number of reads (Fig. 7). Metazoan reads were proportionately more abundant in Bright Bank CTD samples (Cast 8) than in the Slope CTD samples (Casts 14 and 15) (Fig. 7).

Asymptotic rarefaction curves indicated that the sequencing depth was sufficient to capture the diversity in most of the CTD and *Mesobot* samples, and that *Mesobot* samples generally recovered more ASVs than the CTD samples (Fig. 8). The only exception to this pattern was one CTD sample from Cast 15, sampling at 240 m, which detected an unusually high number of ASVs (Fig. 8) although it had slightly less than the average number of sequence reads (40,691 reads) (Supplementary Table 3).

**Fig. 8.**
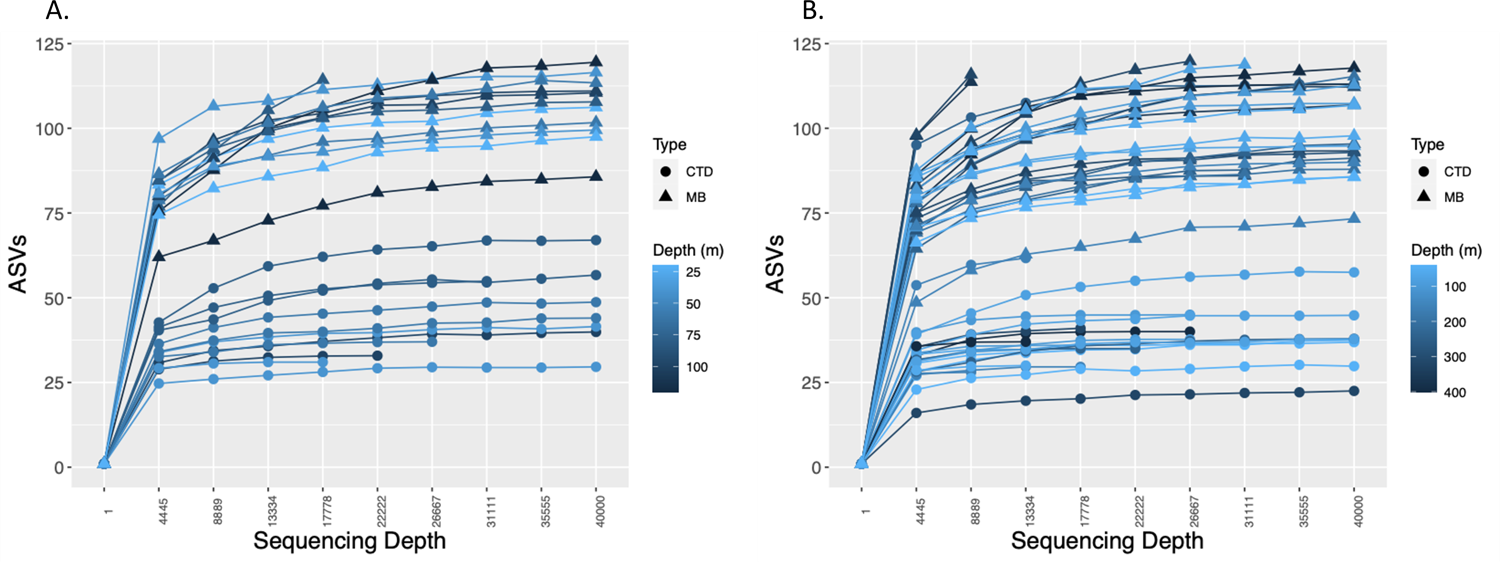
Number of metazoan ASVs in the A) Bright Bank site (MB009 and Cast 8); and B) Slope site (MB011, MB012, Cast 14, and Cast 15). *Mesobot* sampler (MB) samples represent the merged inner and outer filter datasets. Sampling depth is indicated by shade. As some samples had extremely high read counts (>100,000), curves are truncated at 40,0000 in order to visualize all samples, including those with much lower read counts. Total read counts for all samples are in Supplementary Table 3.

### 3.5 Taxonomic composition of the inner and outer sampler filters

The *Mesobot* and CTD samples from both the Bright Bank and Slope sites were comprised of ASVs originating from a wide variety of animal groups (Fig. 9; Fig. 10). Samples were generally dominated by copepod reads (calanoid and cyclopoid) which often comprised the majority of metazoan reads, but ostracods (Halocyprida) and siphonophores were also notably common. Siphonophores occasionally comprised the majority of metazoan reads in some samples, especially in CTD Cast 15 (e.g., at depths 160 m, 320 m, and 400 m at the Slope site). Ostracods were relatively abundant from some samples especially in *Mesobot* deployment MB009 (at the Bright Bank site) at sampling depths 80 m and greater, and in *Mesobot* deployment MB011 (the deep deployment at the Slope site). Very few reads were classified as fish. While the same broad taxonomic groups were generally present among samples, sample biological replicates varied substantially in the relative abundances of taxa (Fig. 9; Fig. 10). Occasionally, it appeared that one taxon would overwhelmingly dominate a particular sample but would be much less common in the corresponding duplicate sample (e.g., siphonophores in samples 320-1 and 400-1 in Cast 15, and in sample 160-1 in MB011; Fig. 9).

**Fig. 9.**
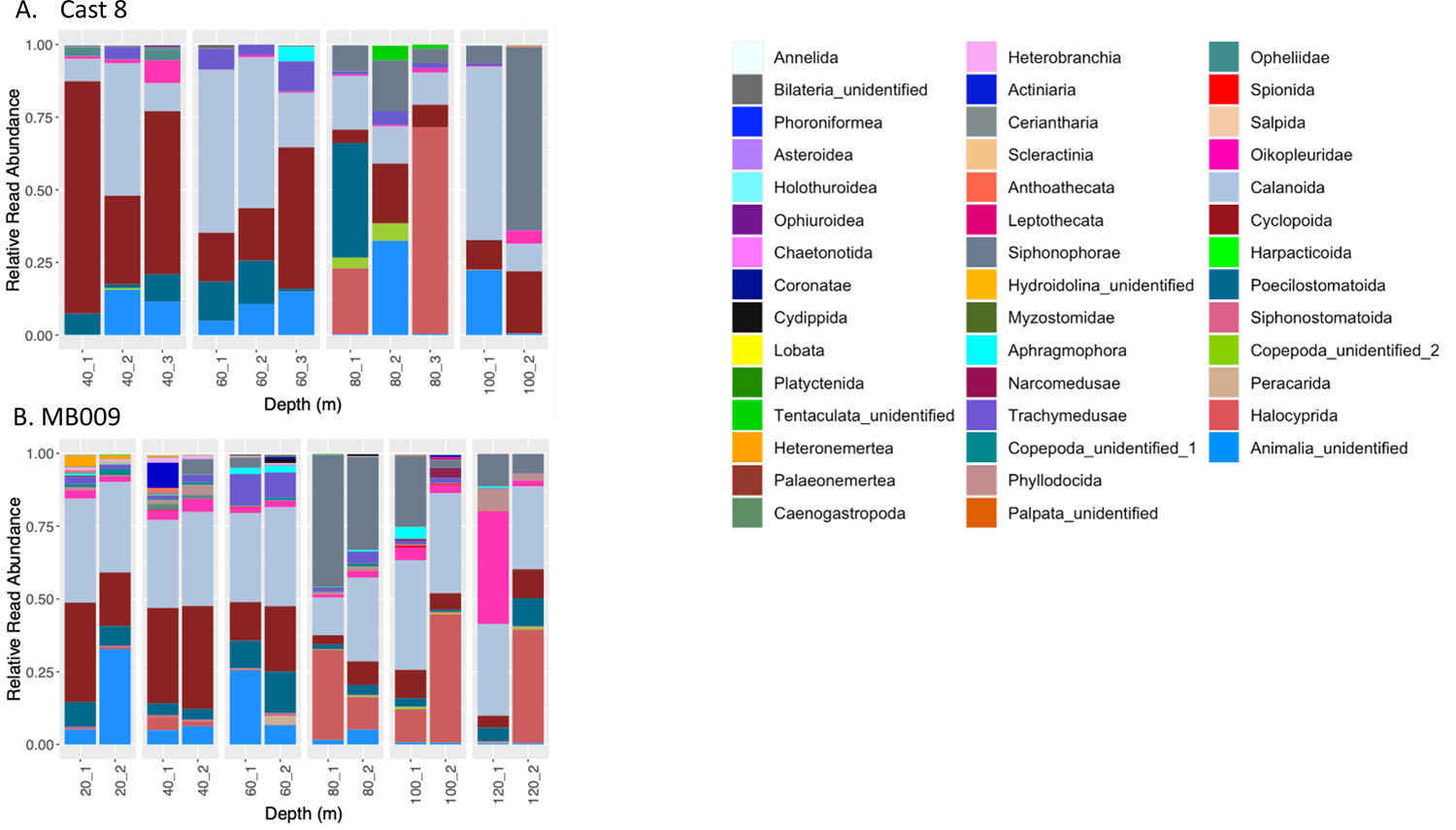
Relative read abundances of Silva level-6 metazoan taxa from the Bright Bank site. A) Cast 8; B) MB009. Only taxa with a summed read frequency across all samples of >500 are shown.

**Fig. 10.**
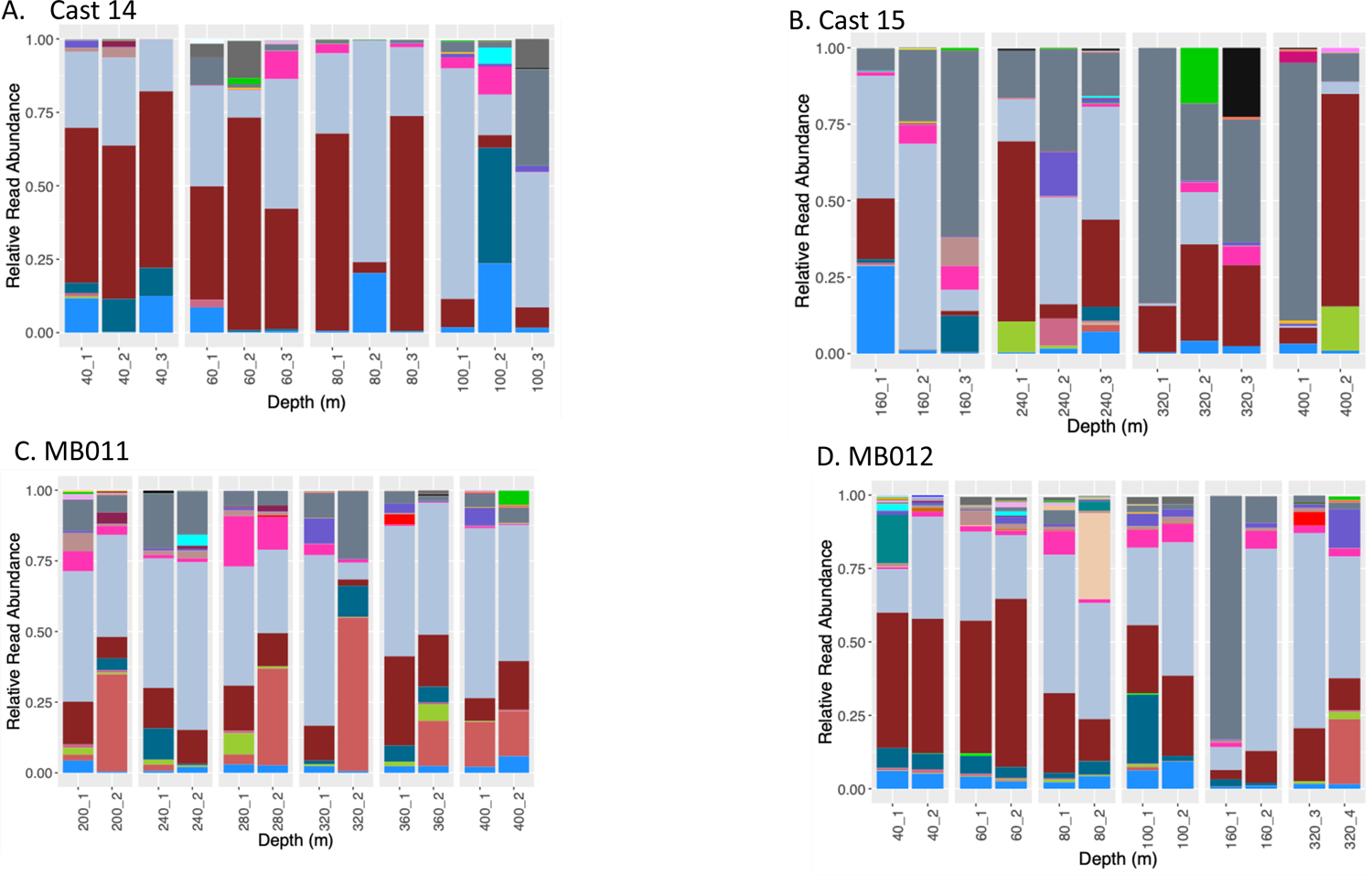
Relative abundances of Silva level-6 taxa from the Slope site. A) Cast 14; B) Cast 15; C) MB011; D) MB012. Only taxa with a summed read frequency of >500 across all samples are shown. Legend is shown in Figure 9.

We compared the Silva level-7 taxa found in samples taken by both methods at a given site and depth. In all but one case, the *Mesobot* samples (duplicates for the site/depth pooled; representing ∼80 – 120 liters of water sampled) detected, on average, 1.66 times more taxa than corresponding CTD samples (triplicates for the site/depth pooled, representing ∼6 liters of water sampled) (Table 3; Appendix 2). There were between 22 – 33 shared taxa (detected in both sampling approaches) depending on the depth, representing on average 36% of all taxa detected at a given depth. There were typically more taxa unique to the *Mesobot* samples (25 – 40) than were unique to the CTD samples (2 – 12; Table 3), representing, on average, 43% (*Mesobot)* and 11% (CTD) of all taxa at a given depth. The one exception was at the Slope site at 240 m depth, where there were 33 taxa detected by both sample types but the CTD samples detected 23 unique taxa and the *Mesobot* detected only 9 unique taxa. One of the CTD replicates from this depth was the same sample noted to have an unusually high number of ASVs (Fig. 8). Also at the Slope site, one depth (320 m) was sampled during two Mesobot deployments (MB011 and MB012) as well as with the CTD. In this case, both *Mesobot* samplings detected more unique taxa than the CTD sampling, and also each *Mesobot* deployment detected several taxa that the other didn’t.

**Table 3.**
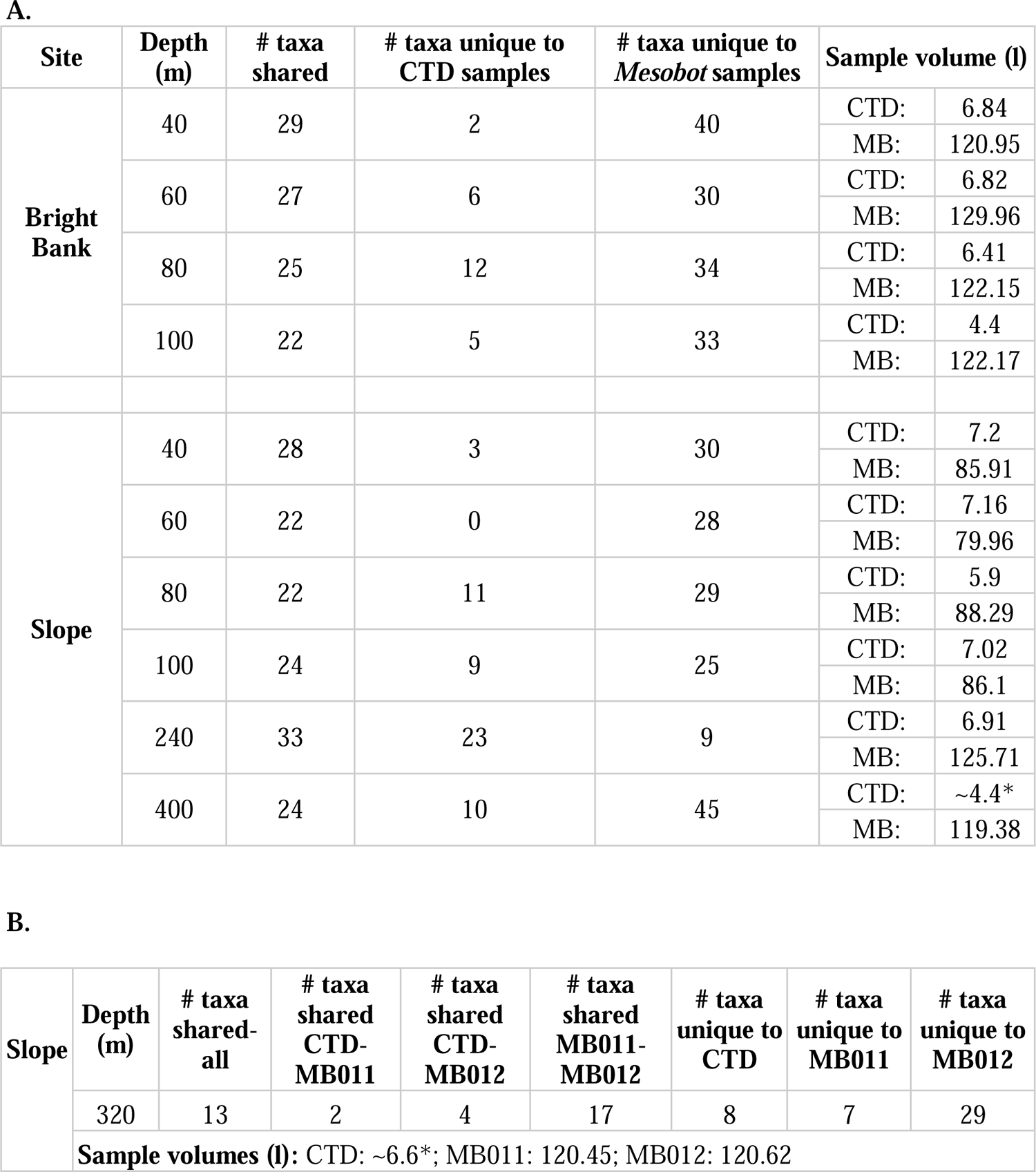
Number of Level-7 taxa at in CTD and *Mesobot* samples from common sites/depths from A) comparisons between 2 sample sets; and B) comparisons between 3 samples sets. *CTD filter volumes not measured; approximations assume 2.2 liters per bottle.

The Bright Bank and Slope datasets were rarefied to their lowest sequencing depths (17,793 and 3,354, respectively) before calculating Bray-Curtis dissimilarities. The nMDS and PERMANOVA analyses indicated structuring relative to sampling depth at the Bright Bank (Fig. 11; sample type: *R^2^* = 0.06688, *p* = 0.013; depth: *R^2^* = 0.51695, *p* = 0.001) and Slope (Fig. 11; sample type: *R^2^* = 0.06181, *p* = 0.001; depth: *R^2^* = 0.41870, *p* = 0.001) sites. Sampling depth had a greater impact than sampling type at the Bright Bank site. These results were supported by functional regressions showed that sampling depth was strongly correlated with the first dimension (MDS1) (Bright Bank: *R^2^* = 0.7551, *p* = 0; Slope: *R^2^* = 0.6218, *p* = 0) but not the second (Bright Bank: *R^2^* = 0.005519, *p* = 0.7439; Slope *R^2^* = 0, p = 0.9905), and no obvious trend with sampling type (Supplementary Fig. 3).

**Fig. 11.**
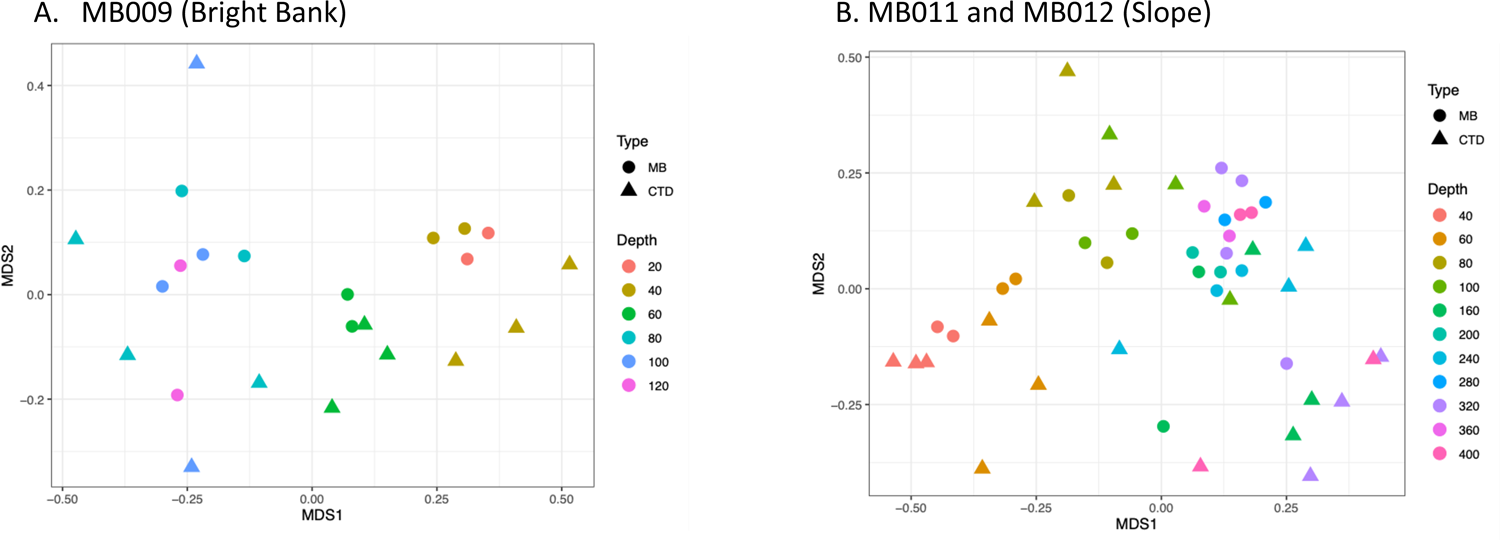
nMDS plots based on Bray-Curtis dissimilarities from the A) MB009 deployment (Bright Bank site), stress = 0.1511615; and B) MB011 and MB012 deployments (Slope site), stress = 0.1815937.

When the inner and outer filters for each *Mesobot* sampler sample were analyzed separately, the relative proportions of the most abundant taxa differed (Fig. 12; Fig. 13). When calculating Bray-Curtis dissimilarities, the dataset was rarefied to 3,438 reads. Four samples from deployment MB009 (1 sample from 20 m, 2 samples frm 40 m, and one sample from 100 m) where the inner filters had read counts below this threshold were excluded. The PERMANOVA results indicated that sampling depth (Bright Bank: *R^2^* = 0.29513, *p* = 0.001; Slope: *R^2^* = 0.15503, *p* = 0.01) had a greater impact than filter type (Bright Bank: *R^2^* = 0.05691, *p* = 0.123; Slope: *R^2^* = 0.04972, *p* = 0.02). This was visualized in the nMDS plot (Fig. 13). Regressions showed that depth was correlated with the first dimension (*R^2^* = 0.8614, *p* = 0) but not the second (*R^2^* = 0.003707, *p* = 0.7932) (Supplementary Fig. 4). In general, gelatinous taxa including siphonophores, trachymedusae, and larvaceans (Oikopleuridae) were more abundant on the inner filters than the outer filters. Out of a total of 181 Silva level-7 (the most highly-resolved level in the Silva classification) taxa, 118 were found on both filter types, 18 on the inner filters only, and 45 on the outer filters only. Notably, there were no crustaceans or fish unique to the inner filters; while there were 7 crustaceans (5 copepods and two eumalacostracans) and two fish unique to the outer filters (Appendix 2). The taxa that were unique to the inner filters were primarily medusozoans, ctenophores, sponges, and polychaetes and other worm-like groups.

**Fig. 12.**
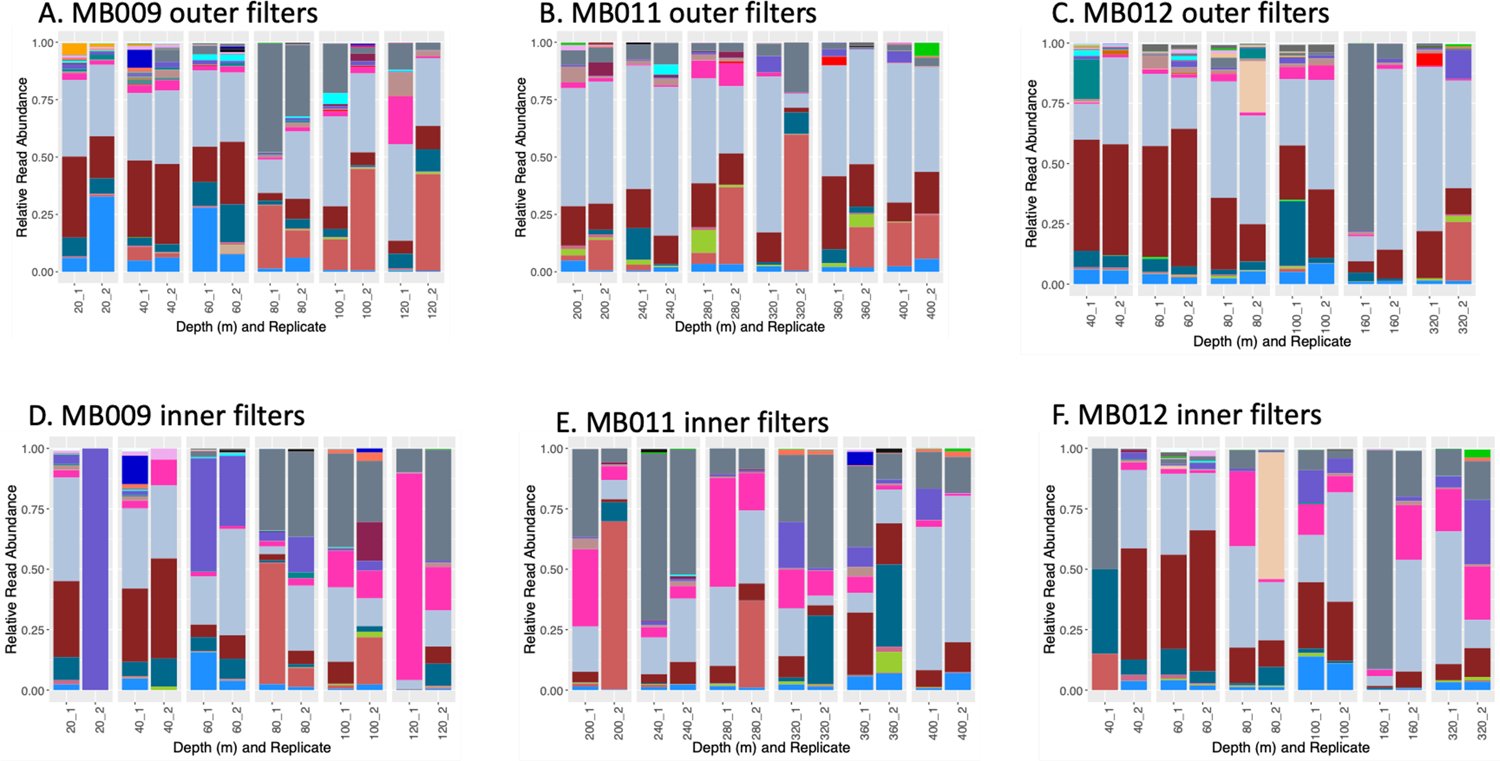
Relative read abundance of level-6 taxa from the outer and inner *Mesobot* filters. A) MB009 (Bright Bank), outer filter; B) MB011 (Slope), outer filter; C) MB012 (Slope), outer filter; D) MB009 (Bright Bank), inner filter; E) MB011 (Slope), inner filter; F) MB012 (Slope), inner filter. Only taxa with a summed read frequency of >500 across all samples are shown. Legend is shown in Figure 9.

**Fig. 13.**
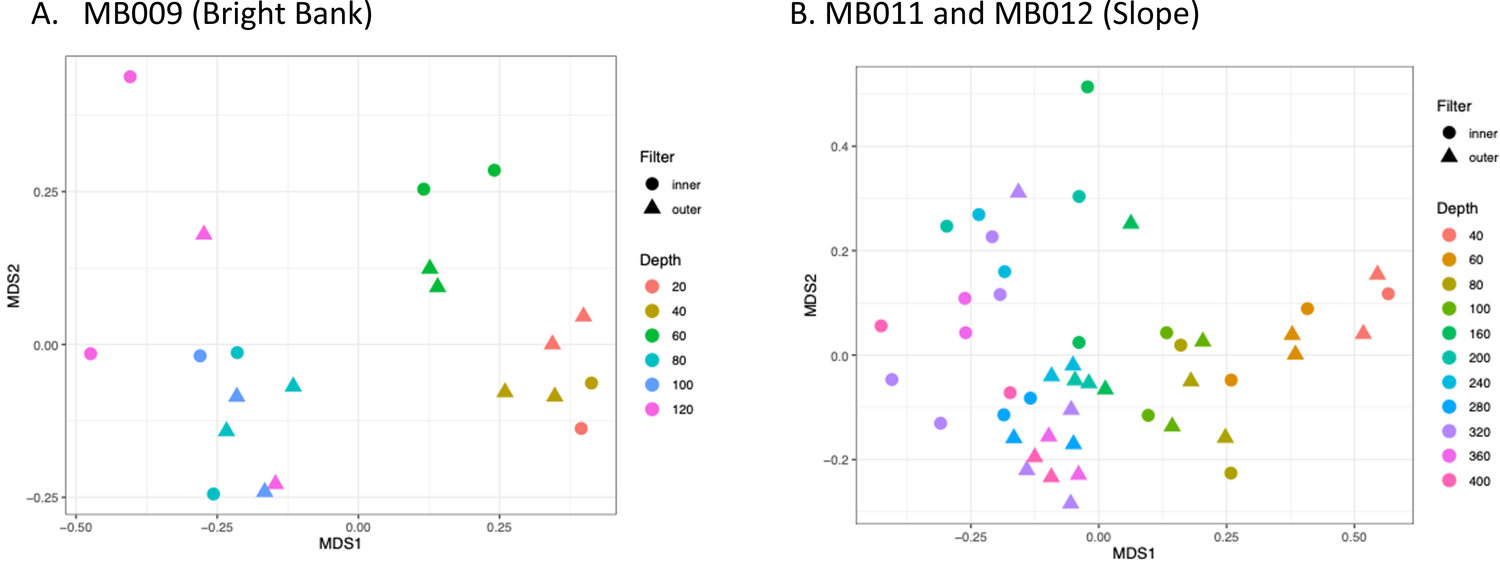
nMDS plots based on Bray-Curtis dissimilarities comparing inner and outer filters and depth from the A) MB009 deployment (Bright Bank site), stress = 0.1436734; and B) the MB011 and MB012 deployments (Slope site), stress = 0.1856701.

## 4 Discussion

We built a large – volume eDNA sampler and successfully deployed it during three dives using *Mesobot* as our sampling platform. Our sampler filtered approximately 20 – 30 times more volume per sample (∼40-60 liters) than our conventionally – obtained CTD samples (∼2 liters). Our hypothesis, that there would be more taxa identified from the large – volume *Mesobot* samples, was supported. We found 66% more taxa in *Mesobot* samples than CTD samples. We also found that the majority of taxa found in the CTD samples were also found in corresponding *Mesobot* samples (78% on average). However, we found that there was also substantial variation between replicates in both the *Mesobot* and CTD sample sets, and despite recovering fewer overall taxa, that the CTD samples did collect unique taxa correspoding to 11% of all taxa sampled at a given depth (compared to 43% taxa sampled only by *Mesobot*). *Mesobot* and CTD sample sets both showed that community composition patterns are strongly associated with depth, thus supporting our hypothesis that despite the differences in taxon detection, the overall community patterns revealed by both methods would be similar.

### 4.1 Sampling volume

While highly variable in both sampling types, our *Mesobot* eDNA capture rate (in terms of the concentration of our extractions as measured by the Qubit fluorometer) was in the same range as for the CTD sampling, after accounting for sample volume and depth. Our study shows a decrease in eDNA with depth that is consistent with previous studies (Govindarajan et al., 2021; McClenaghan et al., 2020). This finding indicates that greater sample volumes are needed for mid and deep water eDNA biodiversity analyses. This is especially true when the focal organisms are animals (as opposed to microbes) – given the small fraction (of metazoan sequence reads we observed in our samples (e.g., <50% in most and <10 % in some) and when the eDNA signal is inhomogeneous.

Sampling approaches and theory are understudied aspects of eDNA protocols (Dickie et al., 2018), and future work should evaluate the optimal sampling volume and strategy as a function of the environment and the biology of target taxa (Mächler et al., 2016). Studies in other environments have similarly demonstrated that increasing sample volumes can improve biodiversity detection (Bessey et al., 2020; Hestetun et al., 2021; Schabacker et al., 2020; Sepulveda et al., 2019). Because our eDNA sampler can efficiently pump a much larger volume than that which can be captured by a single Niskin bottle, it represents a better tool for collecting eDNA at deeper ocean depths (i.e., below ∼100 m). Increasing the sample volume may be especially important for studies in mesopelagic and deeper waters where animal eDNA may be more dilute and when detection of rare taxa is an objective of the study. It is often of interest to obtain vertical profiles in mesopelagic studies, as the vertical dimension is a key axis for environmental variables such as light availability, and for ecological processes such as diel vertical migration. For vertical sampling transects that run from shallow water (e.g., < 100 m, or above the thermocline or DCM) to deep water (e.g., > 100, or below the thermocline or DCM), it may be advantageous to adjust sampling volume with depth (e.g., Laroche et al., 2020).

### 4.2 Filters for large – volume sampling

Filter selection requires special consideration in large-volume eDNA filtering. Previous studies that used larger sample volumes have taken different approaches. Small (submicron) pore size filters which are typically used in eDNA sampling may have slow filtration rates and the filters could become easily clogged (Turner et al., 2014). Some researchers obtain higher sample volumes by utilizing multiple submicron-opening filters (Goldberg et al., 2016; Mächler et al., 2016); but the disadvantages to this approach are the length of time needed to do the filtering and the cost and processing time for multiple filters and their subsequent analyses including additional DNA extractions, PCR, and sequencing. Other studies have utilized larger-pore size filters (Schabacker et al., 2020), but the disadvantage is that taxa that have eDNA predominantly associated with smaller particles could be missed (Sepulveda et al., 2019). Additionally, when large volumes are filtered, it is likely that some intact animals are collected in addition to eDNA. The ideal filter pore size depends on the form of the eDNA of the target taxa; however, eDNA particle sizes are known for only very few taxa (Jo et al., 2019; Moushomi et al., 2019; Turner et al., 2014) Sometimes, a pre-filter to screen out large particles and even whole organisms is used, but using pre-filters may result in the detection of fewer taxa (Djurhuus et al., 2018), unless the pre-filter is also processed.

Our Mini Kleenpak sampler filters had an outer filter with variable-sized pores and an inner filter with 0.2 µm pores and an effective filtration area of 200 cm^2^. For comparison, the Sterivex filters were made of the same material (PES) and the same pore size, but had an order of magnitude smaller filtration area (10 cm^2^). Our sampler outer filters essentially served as a prefilter to the inner filters, and we processed and analyzed both, which added to the effort and cost involved.

The processing included dividing each inner and outer filter into 6 pieces and extracting each, and then pooling and sequencing the inner and outer pieces separately. Thus, each *Mesobot* sample required 12 extractions and 2 pooled PCR reactions per sample for sequencing (versus 1 extraction and 1 pooled PCR reaction for each CTD sample). There is clearly a tradeoff between sample volumes and project cost and effort. As this was the first time that we were aware of that Mini Kleenpak filters were used for eDNA sampling, and the first time that they were used in an offshore marine environment, we elected to process the entirety of the filter area; however, some aspects of our protocols could be refined in the future, as we discuss in section *4.3*.

The outer Mini Kleenpak filters contained a much larger proportion of metazoan sequence reads than the inner filters, indicating a greater retention of animal eDNA on those filters. We observed a reduction in flow rate through all of our Mini Kleenpak filters over time. As the filter pore spaces became reduced or blocked, smaller particles that might have initially passed through the outer filter probably became trapped on the material on the outer filter. Thus, we might expect that eDNA in the form of very small particulates or extracellular DNA could be found on both filters, and that eDNA in the form of larger particulates or even whole animals would be found primarily on the outer filters. We found that most metazoan taxa could be detected from both filter types, but each filter type recovered taxa that the other missed. The taxa found on both filter types included a broad range of animal groups (e.g., medusuzoans, polychaete worms and other worm-like animals, crustaceans, and fish). However, there were many more taxa, originating from a broad range of animal groups, that were unique to the outer filters than to the inner filters. Notably, several crustacean taxa found on the outer filters only but there were no crustaceans unique to the inner filters. The disproportional presence of crustaceans on the outer filters only may suggest their eDNA signal is associated with larger particles, and/or that the outer filters retained zooplankton as well as eDNA.

### 4.3 Logistical considerations

The cost and labor of conducting large volume eDNA sampling and analyses may be higher than for smaller-volume samples as we have noted here and observed elsewhere (Wittwer et al., 2018). From the field perspective, our sampler required about an hour and a half of effort per deployment to prime the pumps, and upon retrieval, the sampler samples could be immediately stored. In contrast, the CTD sampling and processing required more time after retrieval (about four hours of effort per deployment) to filter the same number of samples (12) with around 20 – 30 times less volume per sample. In situations where the number of samples is greater or the sample volumes are larger, the post-retrieval processing time would be even longer, potentially allowing the eDNA signal to decay. Thus, reduction of post-retrieval shipboard processing time is an important advantage of using a sampler with in situ filtration.

Laboratory time and costs are also important to consider. If multiple filters are used to obtain the large volume, the cost of DNA extraction is multiplied. Here, we utilized a single large-area filter, and our DNA extraction protocol necessitated dividing up the filter into pieces for individual extractions. Ideally, only a portion of the filter could be processed and the remainder could be archived (Sepulveda et al., 2019). However, it would need to be shown first that the DNA is distributed evenly throughout the filter, and our data suggest that this is not necessarily the case. If the DNA is not evenly distributed, then by processing only a portion of the filter, the advantages of large volume filtering will be lost. An alternative to this issue would be to develop a DNA extraction protocol that processes the whole filter without having to partition it. For Mini Kleenpak filters, depending on the goal of the study, it might be acceptable to extract only the outer filters which capture the majority of metazoan diversity, although it should be acknowledged that taxa with smaller eDNA particle size distributions could be missed.

Alternatively, the sampler design could be adapted to accommodate other filter types that have only larger openings. Future research with the Mini Kleenpak and other large surface area filters should also explore refinements to the DNA extraction protocol to reduce the cost and labor involved, while preserving the ability to detect a wide range of taxa.

Another relevant sample processing feature that impacts the quantity of taxa detected and should be further explored is the number of PCR replicates in the library preparation step (Ruppert et al., 2019). Increasing the number of PCR replicates increases the number of taxa identified (Ficetola et al., 2015), but also adds to the time and cost of the project. Here, we used duplicate PCRs, but future work should evaluate the benefits of increased replication as this is likely especially important for large volume samples.

### 4.4 General biodiversity observations

Our eDNA analyses from both the *Mesobot* sampler and the CTD sampling revealed a broad range of invertebrate taxa, consistent with what other studies have found with the 18S V9 marker (Blanco-Bercial, 2020; Bucklin et al., 2019; Govindarajan et al., 2021). The paucity of fish reads is also consistent with these other studies, and prior observations that the V9 marker preferentially amplifies taxa other than fish (Sawaya et al., 2019). Sequence reads from crustacean taxa including calanoid and cyclopoid copepods and ostracods were especially abundant in most samples. Siphonophore reads were also common in samples collected at 80 meters and deeper. While the 18S V9 marker detects a wide variety of taxa, it lacks the resolution to identify most taxa to species (Blanco-Bercial, 2020; Bucklin et al., 2016; Wu et al., 2015) and we did not attempt species-level identification in this study. However, future analyses of these samples with other markers could reveal valuable ecological insights on target species. In particular, markers targeting fish such as 12S (e.g., Miya et al., 2015) and anthozoans will be especially relevant for our study site. Additionally, independent methods of characterizing biodiversity such as analyses of net tows and video are important to relate eDNA signatures to community composition (Closek et al., 2019; Govindarajan et al., 2021; Stoeckle et al., 2021). *Mesobot* also has imaging capability (Yoerger et al., 2021) and future studies combining *Mesobot* imaging with our eDNA sampler will reveal further insights into mesophotic and deep water biodiversity.

### 4.5 Biodiversity changes with depth

Despite differences in taxon detection, both of our sampling approaches revealed significant changes in community structure with depth. This is an important finding as it shows that despite the small volumes of water that are sampled, community biodiversity trends can still be detected using conventional CTD/Niskin bottle sampling – which is the most common approach to marine eDNA sampling. Furthermore, despite a myriad of processes that could potentially blur eDNA signatures in oceanic environments – such as particle sinking, ocean currents, vertical mixing, and biologically-mediated transport such as diel vertical migration, our results and other recent studies indicate that eDNA signatures may remain localized. Our finding that eDNA detected diversity changes on the order of 10s of meters in depth are consistent with modeling results that show midwater eDNA signatures remain within 20 meters of their origin in the vertical direction (Allan et al., 2021), and add to a growing body of field evidence from pelagic systems demonstrating that eDNA can detect biodiversity changes with depth (Canals et al., 2021; Easson et al., 2020; Govindarajan et al., 2021).

### 4.6 Variation between replicates

Environmental DNA analyses often show substantial variability between replicates (Beentjes et al., 2019) as we observed here. The optimal number of replicates to include in any eDNA study depends on the study system and goals; however, replication strategies in eDNA studies are inconsistent, and generally not optimized (Dickie et al., 2018). The variation observed here and elsewhere (e.g., Andruszkiewicz et al., 2017; Govindarajan et al., 2021) with CTD sampling suggests that read abundances in individual samples may not be representative of community proportions and that absences of taxa may be false negatives. This variation indicates that eDNA distributions are patchy within a given location or depth, even if eDNA communities are distinguishable between depths.

At our Slope site, the eDNA community at 320 m depth was sampled during both the MB011 and MB012 deployments, as well as with one CTD cast. We found that despite the more intensive sampling effort, each sampling event still recovered unique taxa, and in particular the MB012 sampling event recovered several more taxa (63) than the MB011 sampling event (39) despite similar sample volumes. These differences may be related to eDNA patchiness in the horizontal direction. In mesopelagic depths such as this sampling location, diel vertical migration can create variation in horizontal zooplankton distributions (Chen et al., 2021), which could result in patchy eDNA distributions. More research on the spatial distribution of eDNA in the horizontal dimension of midwater environments would be insightful for optimizing eDNA sampling strategies.

Larger-volume sampling might be expected to lead to more consistent results in biological replicates (which are sampled at the same and location). However, we found that the relative proportions of taxa differed substantially between replicates even in our large-volume *Mesobot* samples. Given the volume of water that we sampled (∼40 - 60 of liters), it is highly likely that small zooplankton were collected along with the eDNA. This possibility is also consistent with our observation of several crustacean taxa unique to the outer filters. If zooplankton are retained on the filters, they would likely be contributing disproportionately to the eDNA reads in that particular sample. Thus, paradoxically, while larger volumes may smooth out variation in eDNA particle distributions, the collection of small zooplankton in addition to particles may introduce a new source of variation. The introduction of a pre-filter to screen out the zooplankton, is not a straightforward solution, as discussed in sections *4.2* and *4.3*.

### 4.7 Autonomous sampling with a robotic platform

The combination of autonomous sampling with robotic platforms and molecular sensing is extremely powerful and has great potential to reveal biological patterns and processes in poorly understood midwater ecosystems (McQuillan and Robidart, 2017). Our sampler successfully obtained large volume eDNA samples from the water column down to 400 m water depth. The sampler was mounted on *Mesobot*, a midwater robot that can operate up to 1000 m depth and track particles and animals whiling utilizing a wide variety of sensors (Yoerger et al., 2021). Our cruise was the second-ever midwater deployment of *Mesobot*. Since the 2019 cruise, the capabilities and operation readiness of the vehicle have expanded. *Mesobot* now carries machine-vision monochrome stereo cameras (Allied Vision G-319B) that enable real-time tracking of midwater targets (Yoerger et al., 2021), a color camera (Sony UMC-SC3A) that provides high-quality color video (HD or 4K) and high-resolution stills (12 MP), and a high-sensitivity radiometer (Oceanic Labs) which can measure downwelling irradiance. Thus, there is great potential to use our sampler with complementary video and environmental data to address a wide variety of midwater hypotheses (Lindsay, 2021). The approach of using *Mesobot* as an eDNA sampling platform opens up a wide range of possible experimental designs that are not possible with traditional CTD sampling, which is limited to vertical casts and the collection of limited volumes of water. Our eDNA sampler could also be integrated on to other platforms, including observational networks (Thorrold et al., 2021).

## 5 Conclusions

We introduced a new eDNA sampler that is capable of filtering large volumes of seawater in situ. We mounted the sampler on the midwater robot *Mesobot* and conducted three successful deployments at two sites in the Flower Garden Banks region of the Gulf of Mexico where we collected samples between 20 and 400 m water depth. We additionally sampled and analyzed eDNA from three CTD casts from the same sites and depths. While both approaches detected biodiversity patterns with depth on the scale of 10s of meters, we found that our large volume samples detected more animal taxa than our conventionally – collected small volume CTD samples. Large-volume sampling could be especially important to consider for mid and deep-water marine environments, and in any environment where eDNA is dilute or patchily – distributed, and when the detection of rare taxa is a goal.

## Acknowledgements

We thank the Scibotics Lab (WHOI) for contributing their midwater oil samplers to the eDNA sampler development, Katie Foley (Lehigh) for assistance with CTD sample processing in the field, and James MacMillan (FGBNMS) for assistance with CTD deployments, Erin Frates (WHOI) for assistance with laboratory sample processing, Peter Wiebe (WHOI) for discussions on data analysis, and Sarah Stover (WHOI) for proof-reading the manuscript. We thank Jessica Labonté and the Ocean & Coastal Studies laboratories of Texas A&M University at Galveston for access to ultrapure water. We also thank the Flower Garden Banks National Marine Sanctuary and the captain and crew of *R/V Manta*.

