## Supplementary Fig. for "Improved biodiversity detection using a large-volume environmental DNA sampler with in situ filtration and implications for marine eDNA sampling strategies"

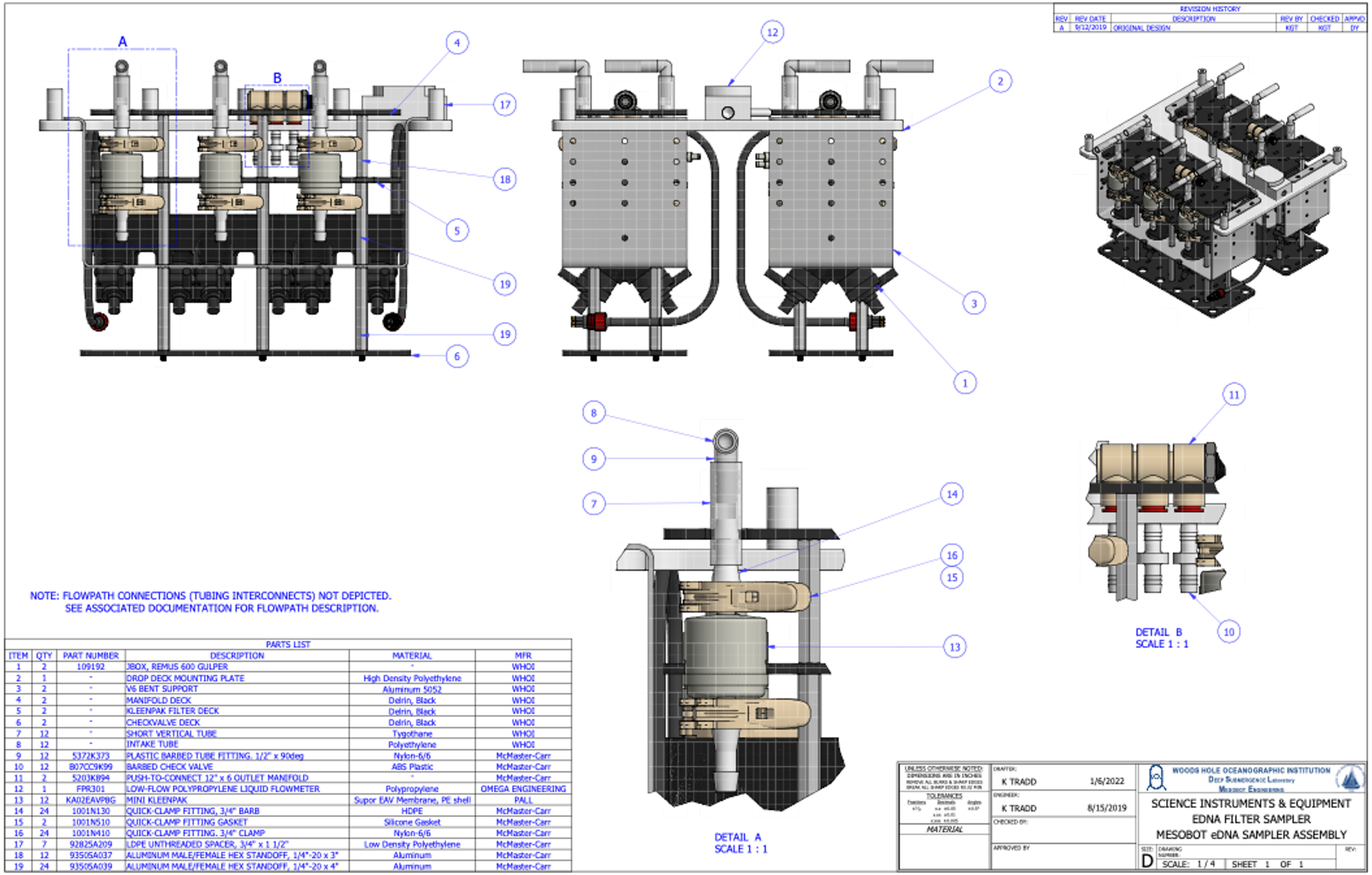


PARTS LIST

| ITEM | QTY | PART NUMBER | DESCRIPTION | MATERIAL | MFR |
|---|---|---|---|---|---|
| 1 | 2 | 109192 | JBOX, REMUS 600 GULPER | - | WHOI |
| 2 | 1 | - | DROP DECK MOUNTING PLATE | High Density Polyethylene | WHOI |
| 3 | 2 | - | V6 BENT SUPPORT | Aluminum 5052 | WHOI |
| 4 | 2 | - | MANIFOLD DECK | Delrin, Black | WHOI |
| 5 | 2 | - | KLEENPAK FILTER DECK | Delrin, Black | WHOI |
| 6 | 2 | - | CHECKVALVE DECK | Delrin, Black | WHOI |
| 7 | 12 | - | SHORT VERTICAL TUBE | Tygothane | WHOI |
| 8 | 12 | - | INTAKE TUBE | Polyethylene | WHOI |
| 9 | 12 | 5372K373 | PLASTIC BARBED TUBE FITTING. 1/2" x 90deg | Nylon-6/6 | McMaster-Carr |
| 10 | 12 | B07CC9K99 | BARBED CHECK VALVE | ABS Plastic | McMaster-Carr |
| 11 | 2 | 5203K894 | PUSH-TO-CONNECT 12" x 6 OUTLET MANIFOLD | - | McMaster-Carr |
| 12 | 1 | FPR301 | LOW-FLOW POLYPROPYLENE LIQUID FLOWMETER | Polypropylene | OMEGA ENGINEERING |
| 13 | 12 | KA02EAVP8G | MINI KLEENPAK | Supor EAV Membrane, PE shell | PALL |
| 14 | 24 | 1001N130 | QUICK-CLAMP FITTING, 3/4" BARB | HDPE | McMaster-Carr |
| 15 | 2 | 1001N510 | QUICK-CLAMP FITTING GASKET | Silicone Gasket | McMaster-Carr |
| 16 | 24 | 1001N410 | QUICK-CLAMP FITTING. 3/4" CLAMP | Nylon-6/6 | McMaster-Carr |
| 17 | 7 | 92825A209 | LDPE UNTHREADED SPACER, 3/4" x 1 1/2" | Low Density Polyethylene | McMaster-Carr |
| 18 | 12 | 93505A037 | ALUMINUM MALE/FEMALE HEX STANDOFF, 1/4"-20 x 3" | Aluminum | McMaster-Carr |
| 19 | 24 | 93505A039 | ALUMINUM MALE/FEMALE HEX STANDOFF, 1/4"-20 x 4" | Aluminum | McMaster-Carr |

**Supplementary Fig. 1.** eDNA sampler design details.

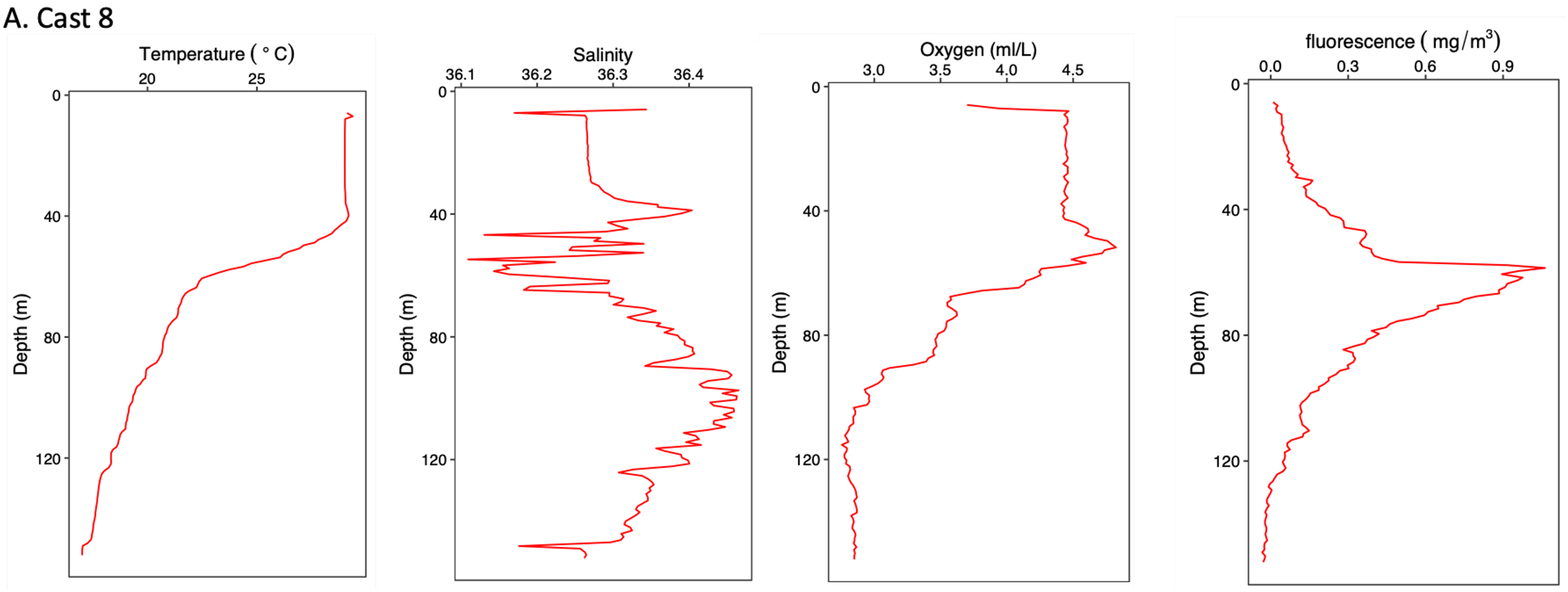


**Supplementary Fig. 2.** CTD data from the A) Bright Bank site - CTD Cast 8; B) Slope site – CTD Cast 14; and C) Slope site – CTD Cast 15.

B. Cast 14

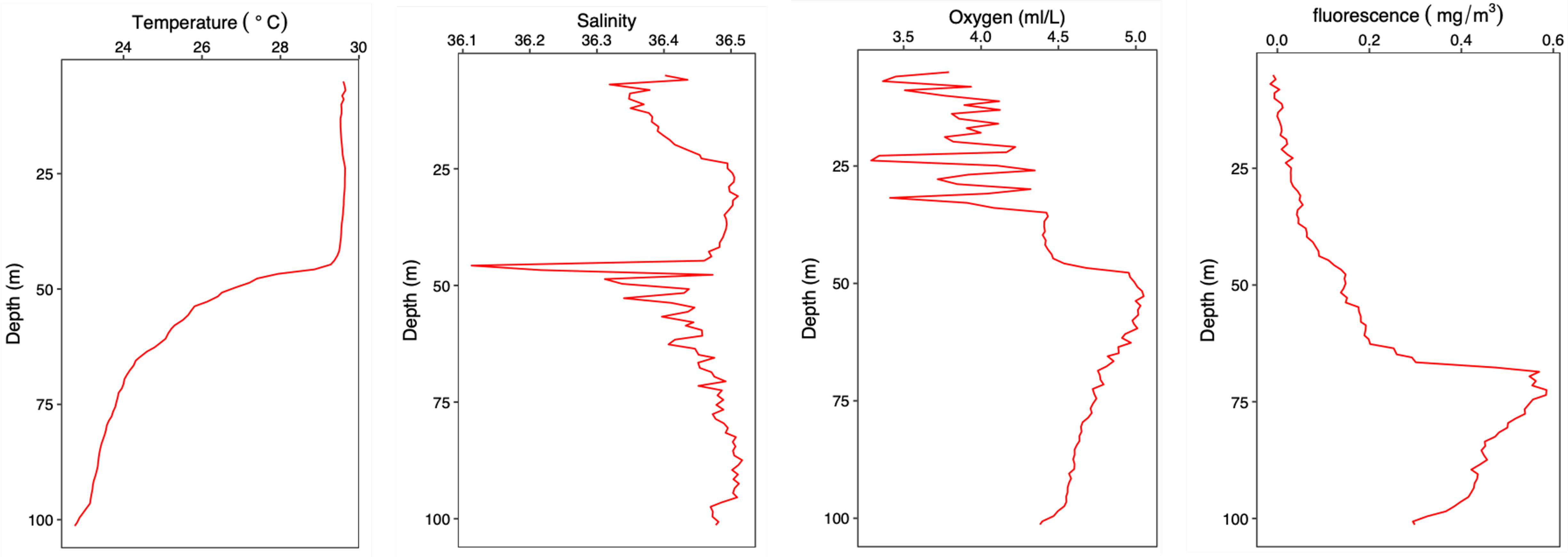
Temperature ( ° C)
24
26
28
30
Depth (m)
25
50
75
100
Salinity
36.1
36.2
36.3
36.4
36.5
Depth (m)
25
50
75
100
Oxygen (ml/L)
3.5
4.0
4.5
5.0
Depth (m)
25
50
75
100
fluorescence ( mg/m3)
0.0
0.2
0.4
0.6
Depth (m)
25
50
75
100

### C. Cast 15

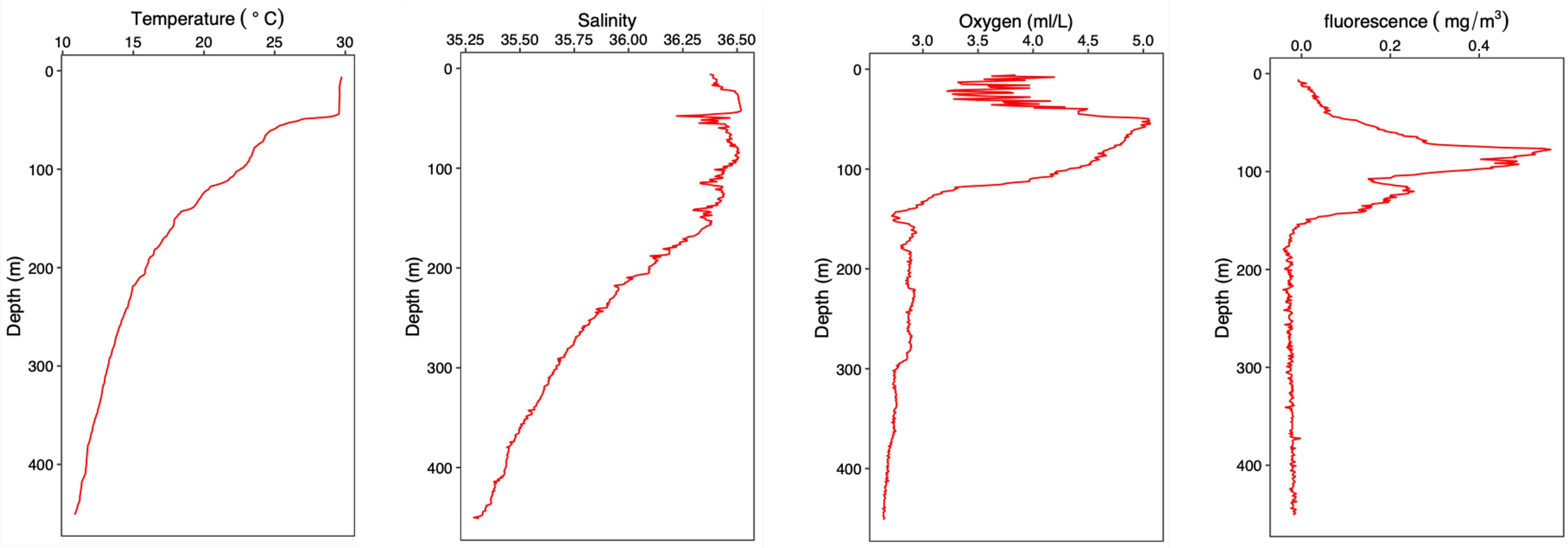
Temperature ( ° C)
10
15
20
25
30
0
100
200
300
400
Depth (m)
Salinity
35.25
35.50
35.75
36.00
36.25
36.50
0
100
200
300
400
Depth (m)
Oxygen (ml/L)
3.0
3.5
4.0
4.5
5.0
0
100
200
300
400
Depth (m)
fluorescence ( mg/m³)
0.0
0.2
0.4
0
100
200
300
400
Depth (m)

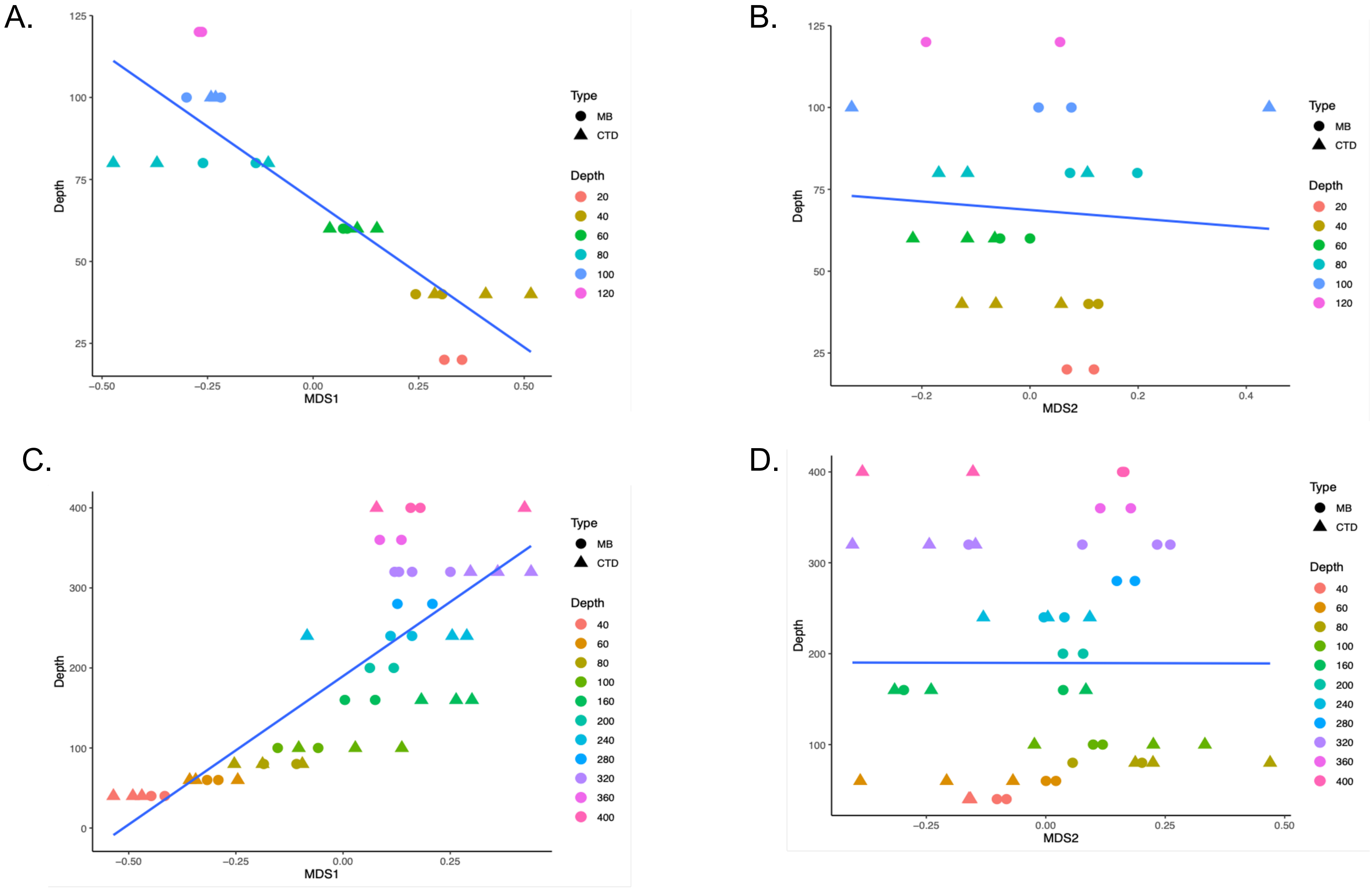


**Supplementary Fig. 3**. Regressions of depth against nMDS axes from Figure 10. A) Bright Bank, Depth vs. MDS1; B) Bright Bank, Depth vs. MDS2; C) Slope, Depth vs. MDS1; D) Slope, Depth vs. MDS2.

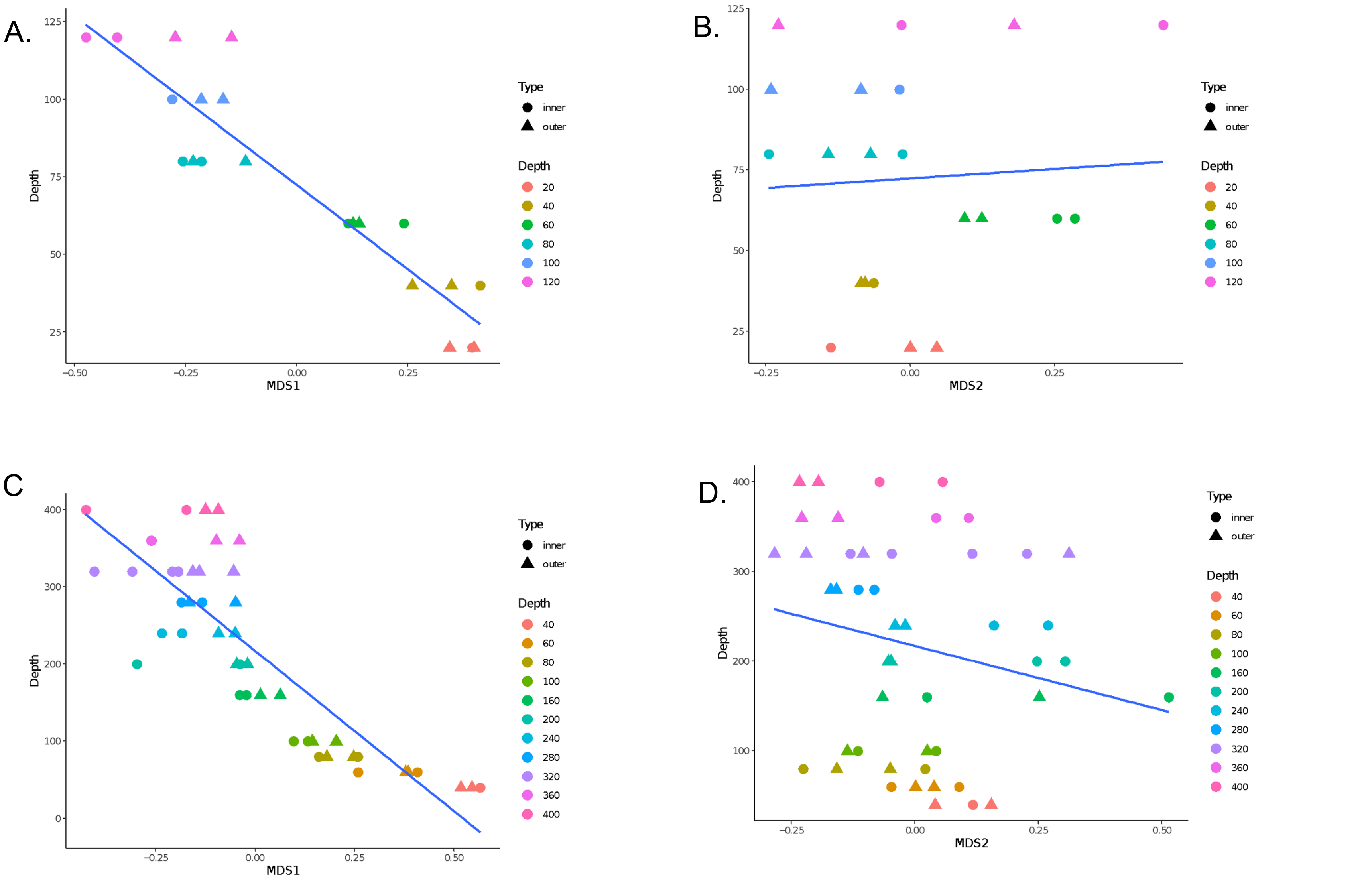


**Supplementary Fig. 4**. Regressions of depth against nMDS axes from Figure 12. A) Bright Bank, Depth vs. MDS1; B) Bright Bank, Depth vs. MDS2; C) Slope, Depth vs. MDS1; D) Slope, Depth vs. MDS2.
