## Supplementary Table for "Improved biodiversity detection using a large-volume environmental DNA sampler with in situ filtration and implications for marine eDNA sampling strategies"

**Supplementary Table 1.** *Mesobot* samples and associated collection information.

| **Dive #** | **Pump** | **Depth (m)** | **Date** | **Station** | **Latitude** | **Longitude** | **Volume (L)** |
| --- | --- | --- | --- | --- | --- | --- | --- |
| MB009 | C1 | 120 | 20190925 | Bright Bank | 27.8485 | -93.2576 | 65.82 |
| MB009 | D1 | 120 | 20190925 | Bright Bank | 27.8485 | -93.2576 | 63.21 |
| MB009 | C2 | 100 | 20190925 | Bright Bank | 27.8485 | -93.2576 | 65.74 |
| MB009 | D2 | 100 | 20190925 | Bright Bank | 27.8485 | -93.2576 | 56.43 |
| MB009 | C3 | 80 | 20190925 | Bright Bank | 27.8485 | -93.2576 | 61.67 |
| MB009 | D3 | 80 | 20190925 | Bright Bank | 27.8485 | -93.2576 | 60.48 |
| MB009 | C4 | 60 | 20190925 | Bright Bank | 27.8485 | -93.2576 | 67.67 |
| MB009 | D4 | 60 | 20190925 | Bright Bank | 27.8485 | -93.2576 | 62.29 |
| MB009 | C5 | 40 | 20190925 | Bright Bank | 27.8485 | -93.2576 | 63.28 |
| MB009 | D5 | 40 | 20190925 | Bright Bank | 27.8485 | -93.2576 | 57.67 |
| MB009 | C6 | 20 | 20190925 | Bright Bank | 27.8485 | -93.2576 | 65.57 |
| MB009 | D6 | 20 | 20190925 | Bright Bank | 27.8485 | -93.2576 | 59.73 |
| MB009 | control | N/A | 20190925 | Bright Bank | 27.8485 | -93.2576 | N/A |
| MB011 | A1 | 400 | 20190926 | Slope | 27.53905 | -93.34029 | 60.94 |
| MB011 | B1 | 400 | 20190926 | Slope | 27.53905 | -93.34029 | 58.44 |
| MB011 | A2 | 360 | 20190926 | Slope | 27.53905 | -93.34029 | 60.5 |
| MB011 | B2 | 360 | 20190926 | Slope | 27.53905 | -93.34029 | 62.87 |
| MB011 | A3 | 320 | 20190926 | Slope | 27.53905 | -93.34029 | 61.77 |
| MB011 | B3 | 320 | 20190926 | Slope | 27.53905 | -93.34029 | 58.68 |
| MB011 | A4 | 280 | 20190926 | Slope | 27.53905 | -93.34029 | 60.94 |
| MB011 | B4 | 280 | 20190926 | Slope | 27.53905 | -93.34029 | 59.78 |
| MB011 | A5 | 240 | 20190926 | Slope | 27.53905 | -93.34029 | 62.51 |
| MB011 | B5 | 240 | 20190926 | Slope | 27.53905 | -93.34029 | 62.66 |
| MB011 | A6 | 200 | 20190926 | Slope | 27.53905 | -93.34029 | 61.19 |
| MB011 | B6 | 200 | 20190926 | Slope | 27.53905 | -93.34029 | 65.22 |
| MB011 | control | N/A | 20190926 | Slope | 27.53905 | -93.34029 | N/A |
| MB012 | C1 | 320 | 20190926-27 | Slope | 27.53905 | -93.34029 | 59.88 |
| MB012 | D1 | 320 | 20190926-27 | Slope | 27.53905 | -93.34029 | 60.74 |
| MB012 | C2 | 160 | 20190926-27 | Slope | 27.53905 | -93.34029 | 41.79 |
| MB012 | D2 | 160 | 20190926-27 | Slope | 27.53905 | -93.34029 | 38.73 |
| MB012 | C3 | 100 | 20190926-27 | Slope | 27.53905 | -93.34029 | 41.23 |
| MB012 | D3 | 100 | 20190926-27 | Slope | 27.53905 | -93.34029 | 44.87 |
| MB012 | C4 | 80 | 20190926-27 | Slope | 27.53905 | -93.34029 | 46.12 |
| MB012 | D4 | 80 | 20190926-27 | Slope | 27.53905 | -93.34029 | 42.17 |
| MB012 | C5 | 60 | 20190926-27 | Slope | 27.53905 | -93.34029 | 41.56 |
| MB012 | D5 | 60 | 20190926-27 | Slope | 27.53905 | -93.34029 | 38.4 |
| MB012 | C6 | 40 | 20190926-27 | Slope | 27.53905 | -93.34029 | 44.61 |
| MB012 | D6 | 40 | 20190926-27 | Slope | 27.53905 | -93.34029 | 41.3 |
| MB012 | control | N/A | 20190926-27 | Slope | 27.53905 | -93.34029 | N/A |

**Supplementary Table 2.** CTD samples and associated collection information.

| **Cast** | **Depth (m)** | **Date** | **Station** | **Latitude** | **Longitude** | **Volume (L)** |
| --- | --- | --- | --- | --- | --- | --- |
| 8 | 100 | 20190925 | Bright Bank | 27.84239 | -93.26850 | 2.15 |
| 8 | 100 | 20190925 | Bright Bank | 27.84239 | -93.26850 | 2.25 |
| 8 | 80 | 20190925 | Bright Bank | 27.84239 | -93.26850 | 2.16 |
| 8 | 80 | 20190925 | Bright Bank | 27.84239 | -93.26850 | 2.06 |
| 8 | 80 | 20190925 | Bright Bank | 27.84239 | -93.26850 | 2.19 |
| 8 | 60 | 20190925 | Bright Bank | 27.84239 | -93.26850 | 2.3 |
| 8 | 60 | 20190925 | Bright Bank | 27.84239 | -93.26850 | 2.26 |
| 8 | 60 | 20190925 | Bright Bank | 27.84239 | -93.26850 | 2.26 |
| 8 | 40 | 20190925 | Bright Bank | 27.84239 | -93.26850 | 2.3 |
| 8 | 40 | 20190925 | Bright Bank | 27.84239 | -93.26850 | 2.3 |
| 8 | 40 | 20190925 | Bright Bank | 27.84239 | -93.26850 | 2.24 |
| 8 | N/A | 20190925 | CTD08 filtration blank | N/A | N/A | 2.4 |
| 14 | 100 | 2019026 | Slope | 27.54012 | -93.35027 | 2 |
| 14 | 100 | 2019026 | Slope | 27.54012 | -93.35027 | 2.32 |
| 14 | 100 | 2019026 | Slope | 27.54012 | -93.35027 | 2.7 |
| 14 | 80 | 2019026 | Slope | 27.54012 | -93.35027 | 2.24 |
| 14 | 80 | 2019026 | Slope | 27.54012 | -93.35027 | 1.5 |
| 14 | 80 | 2019026 | Slope | 27.54012 | -93.35027 | 2.16 |
| 14 | 60 | 2019026 | Slope | 27.54012 | -93.35027 | 2.42 |
| 14 | 60 | 2019026 | Slope | 27.54012 | -93.35027 | 2.34 |
| 14 | 60 | 2019026 | Slope | 27.54012 | -93.35027 | 2.4 |
| 14 | 40 | 2019026 | Slope | 27.54012 | -93.35027 | 2.4 |
| 14 | 40 | 2019026 | Slope | 27.54012 | -93.35027 | 2.4 |
| 14 | 40 | 2019026 | Slope | 27.54012 | -93.35027 | 2.4 |
| N/A | N/A | 2019026 | CTD14 filtration blank | N/A | N/A | 2.41 |
| 15 | 400 | 20190926 | Slope | 27.54607 | -93.38611 | Not taken |
| 15 | 400 | 20190926 | Slope | 27.54607 | -93.38611 | Not taken |
| 15 | 320 | 20190926 | Slope | 27.54607 | -93.38611 | Not taken |
| 15 | 320 | 20190926 | Slope | 27.54607 | -93.38611 | 2.26 |
| 15 | 320 | 20190926 | Slope | 27.54607 | -93.38611 | 2.24 |
| 15 | 240 | 20190926 | Slope | 27.54607 | -93.38611 | 2.4 |
| 15 | 240 | 20190926 | Slope | 27.54607 | -93.38611 | 2.13 |
| 15 | 240 | 20190926 | Slope | 27.54607 | -93.38611 | 2.38 |
| 15 | 160 | 20190926 | Slope | 27.54607 | -93.38611 | 2.4 |
| 15 | 160 | 20190926 | Slope | 27.54607 | -93.38611 | 2.15 |
| 15 | 160 | 20190926 | Slope | 27.54607 | -93.38611 | 2.05 |
| N/A | N/A | 20190926 | CTD15 filtration blank | N/A | N/A | N/A |

**Supplementary Table 3.** Number of metazoan sequence reads per sample.

| **CTD Samples** | |  | **Mesobot Samples** | | | | |
| --- | --- | --- | --- | --- | --- | --- | --- |
| **CTD samples** | **# reads** |  | **Mesobot samples** | **# reads on inner filter** | **# reads on outer filter** | **total # of reads** | **% of reads on outer filter** |
| CTD8-01 | 61616 |  | MB009C1 | 25492 | 66775 | 92267 | 72.37 |
| CTD8-02 | 23593 |  | MB009C2 | 7895 | 37579 | 45474 | 82.64 |
| CTD8-04 | 77219 |  | MB009C3 | 17442 | 101161 | 118603 | 85.29 |
| CTD8-05 | 34607 |  | MB009C4 | 16923 | 66719 | 83642 | 79.77 |
| CTD8-06 | 50769 |  | MB009C5 | 23747 | 79747 | 103494 | 77.05 |
| CTD8-07 | 29863 |  | MB009C6 | 23251 | 65676 | 88927 | 73.85 |
| CTD8-08 | 88488 |  | MB009D1 | 4437 | 55754 | 60191 | 92.63 |
| CTD8-09 | 66873 |  | MB009D2 | 174 | 49447 | 49621 | 99.65 |
| CTD8-10 | 17793 |  | MB009D3 | 11934 | 45225 | 57159 | 79.12 |
| CTD8-11 | 76795 |  | MB009D4 | 28074 | 73882 | 101956 | 72.46 |
| CTD8-12 | 81148 |  | MB009D5 | 981 | 95273 | 96254 | 98.98 |
| CTD14-01 | 19495 |  | MB009D6 | 3 | 207391 | 207394 | 100.00 |
| CTD14-02 | 27961 |  | MB011A1 | 8933 | 42677 | 51610 | 82.69 |
| CTD14-03 | 29814 |  | MB011A2 | 3438 | 55921 | 59359 | 94.21 |
| CTD14-04 | 52949 |  | MB011A3 | 9094 | 49199 | 58293 | 84.40 |
| CTD14-05 | 67501 |  | MB011A4 | 8682 | 23530 | 32212 | 73.05 |
| CTD14-06 | 64947 |  | MB011A5 | 18821 | 72955 | 91776 | 79.49 |
| CTD14-07 | 14254 |  | MB011A6 | 10624 | 56247 | 66871 | 84.11 |
| CTD14-08 | 5638 |  | MB011B1 | 6900 | 33754 | 40654 | 83.03 |
| CTD14-09 | 20614 |  | MB011B2 | 8812 | 89392 | 98204 | 91.03 |
| CTD14-10 | 99996 |  | MB011B3 | 10265 | 112010 | 122275 | 91.60 |
| CTD14-11 | 59236 |  | MB011B4 | 15301 | 38049 | 53350 | 71.32 |
| CTD14-12 | 99783 |  | MB011B5 | 15558 | 94349 | 109907 | 85.84 |
| CTD15-02 | 14772 |  | MB011B6 | 38261 | 63594 | 101855 | 62.44 |
| CTD15-03 | 27522 |  | MB012C1 | 12113 | 88283 | 100396 | 87.93 |
| CTD15-04 | 82223 |  | MB012C2 | 68149 | 103357 | 171506 | 60.26 |
| CTD15-05 | 18099 |  | MB012C3 | 11983 | 74706 | 86689 | 86.18 |
| CTD15-06 | 3354 |  | MB012C4 | 10113 | 47222 | 57335 | 82.36 |
| CTD15-07 | 44167 |  | MB012C5 | 12340 | 89659 | 101999 | 87.90 |
| CTD15-08 | 22885 |  | MB012C6 | 143 | 136461 | 136604 | 99.90 |
| CTD15-09 | 40691 |  | MB012D1 | 8645 | 80283 | 88928 | 90.28 |
| CTD15-10 | 17275 |  | MB012D2 | 9516 | 35270 | 44786 | 78.75 |
| CTD15-11 | 26160 |  | MB012D3 | 18665 | 46226 | 64891 | 71.24 |
| CTD15-12 | 9277 |  | MB012D4 | 46081 | 129367 | 175448 | 73.74 |
|  |  |  | MB012D5 | 29377 | 88567 | 117944 | 75.09 |
|  |  |  | MB012D6 | 40079 | 104710 | 144789 | 72.32 |
