## Appendix 1 for "Improved biodiversity detection using a large-volume environmental DNA sampler with in situ filtration and implications for marine eDNA sampling strategies"

Two control samples were mis-processed during the MiSeq runs and produced a large number of sequencing reads: after the DADA2 and quality control steps, there were119,646 reads in the CTD control (for Cast 16; which was part of the larger Bright Bank Survey but not analyzed here) and 116,805 reads in one of the PCR no template controls. These same controls had no detectable DNA after the library preparation PCRs and produced extremely few reads during the preliminary Miniseq run. We re-sequenced these two controls, plus four additional environmental samples (2 *Mesobot* and 2 CTD samples), across 3 new Miseq runs with the same target sequencing depth as in the original MiSeq runs. As expected, the suspect controls produced very few reads in the second sequencing run series: after the dada2 and quality control steps, there were 0 reads in the CTD control and 112 reads in the PCR no template control. The taxonomic composition of the data from the re-sequenced environmental samples was nearly identical to the composition of the data from samples in the original run series (Supplementary Figure 1). There was no evidence that these or any other environmental samples or controls were compromised in the original MiSeq runs. We therefore proceeded with our data from the original sequencing run series, with the substitution of the two mis-processed control data with the control data from the re-sequenced run.

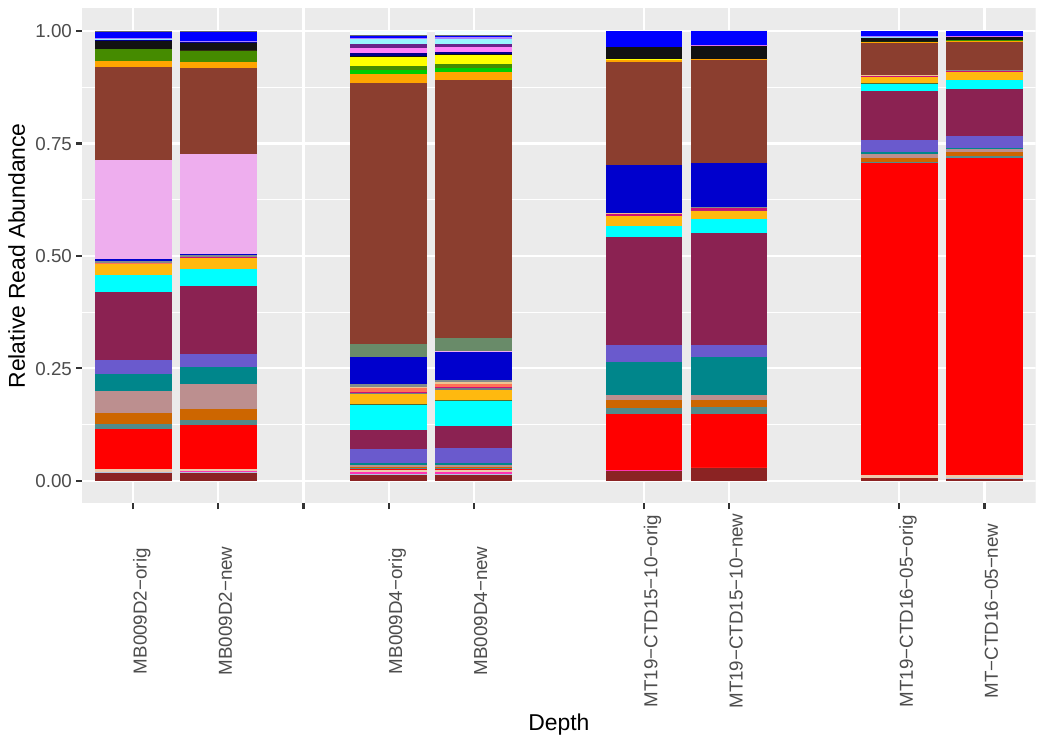
